# Quantitative sub-cellular acyl-CoA analysis reveals distinct nuclear regulation

**DOI:** 10.1101/2020.07.30.229468

**Authors:** Sophie Trefely, Katharina Huber, Joyce Liu, Michael Noji, Stephanie Stransky, Jay Singh, Mary T. Doan, Claudia D. Lovell, Eliana von Krusenstiern, Helen Jiang, Anna Bostwick, Hannah L. Pepper, Luke Izzo, Steven Zhao, Jimmy P. Xu, Kenneth C. Bedi, J. Eduardo Rame, Juliane G. Bogner-Strauss, Clementina Mesaros, Simone Sidoli, Kathryn E. Wellen, Nathaniel W. Snyder

**Author notes:** Co-corresponding authors, **Corresponding authors address:** Nathaniel W. Snyder, Post: 455 Medical Education Research Building, 3500 N Broad St, Philadelphia, PA 19140, USA, Kathryn E. Wellen, Post: 421 Curie Blvd, 653 BRBII/III, Philadelphia, PA 19104-6160, USA.

## Abstract

Quantitative sub-cellular metabolomic measurements can yield crucial insights into the roles of metabolites in cellular processes, but are subject to multiple confounding factors. We developed <u>S</u>table <u>I</u>sotope <u>L</u>abeling of <u>E</u>ssential nutrients in cell <u>C</u>ulture – <u>S</u>ub-cellular <u>F</u>ractionation (SILEC-SF), which uses isotope labeled internal standard controls that are present throughout fractionation and processing to quantify acyl-Coenzyme A thioesters in sub-cellular compartments by liquid chromatography-mass spectrometry. We tested SILEC-SF in a range of sample types and examined the compartmentalized responses to oxygen tension, cellular differentiation, and nutrient availability. Application of SILEC-SF to the challenging analysis of the nuclear compartment revealed a nuclear acyl-CoA profile distinct from that of the cytosol, with notable nuclear enrichment of propionyl-CoA. Using isotope tracing we identified the branched chain amino acid (BCAA) isoleucine as a major metabolic source of nuclear propionyl-CoA and histone propionylation, thus revealing a new mechanism of crosstalk between metabolism and the epigenome.

## Introduction

Quantitative measurements of metabolites can yield crucial insights into their roles in cellular processes and nutrient sensing mechanisms. However, metabolism is highly compartmentalized within cells, and the sub-cellular distribution of metabolites determines their use (Boon et al., 2020; Trefely et al., 2020). Whole cell analyses often provide insufficient information on compartment-specific metabolism, particularly when the compartment of interest accounts for a small fraction of whole cell levels of a given metabolite. Indeed, recent advances in fractionation methodologies for compartment-specific metabolite quantitation by mass spectrometry have yielded important biological insights (Chen et al., 2017; Wyant et al., 2017). Yet, distinct challenges in measuring metabolites in subcellular compartments remain. In particular, sub-cellular fractionation is inherently disruptive and metabolic analysis after fractionation is subject to multiple confounding factors (Dietz, 2017; Lu et al., 2017; Trefely et al., 2019).

Acyl-Coenzyme A thioesters (acyl-CoAs) are a family of metabolites that are involved in multiple metabolic pathways (Trefely et al., 2020). Acetyl-CoA, a prototypical acyl-CoA, is the product of several catabolic processes in the mitochondria, whilst in the cytosol it is the primary substrate for anabolic processes including de-novo fatty acid and cholesterol synthesis. Acetyl-CoA is the acyl-donor for acetylation in all compartments. In the nucleus, acetyl-CoA is a substrate for epigenetic regulation via histone acetylation, and histone acetylation levels have been correlated with cellular acetyl-CoA levels (Cai et al., 2011; Cluntun et al., 2015; Lee et al., 2014).

Metabolite availability in the nucleus can link environmental cues with nuclear gene regulation via nutrient-sensitive chromatin modifications (Campbell and Wellen, 2018; Dai et al.; Ryan et al., 2019). The nucleus and cytosol have been generally considered a continuous metabolic compartment since the nuclear pores are permeable to small molecule metabolites such as acetyl-CoA. Nevertheless, whole cell acetyl-CoA abundance and histone acetylation levels can be uncoupled (e.g., in cells deficient in the acetyl-CoA generating enzyme ATP citrate lyase (ACLY)(Zhao et al., 2016)), and several recent studies have provided evidence of distinct regulation of acetyl-CoA production within the nucleus (Li et al., 2017; Mews et al., 2017; Sivanand et al., 2017, 2018; Sun et al., 2019; Sutendra et al., 2014). Other histone modifications including succinylation, malonylation, propionylation, crotonoylation and butyrylation are also correlated with the cellular abundance of their respective acyl-CoAs (Simithy et al., 2017), but the subcellular distribution and regulation of these metabolites remain poorly understood. Rigorous mass spectrometry-based methodologies for determination of nuclear metabolite levels have not been reported, though such approaches are needed to advance mechanistic understanding of the links between metabolism and chromatin modification.

We developed <u>S</u>table <u>I</u>sotope <u>L</u>abeling of <u>E</u>ssential nutrients in cell <u>C</u>ulture – <u>S</u>ub-cellular <u>F</u>ractionation (SILEC-SF) to quantitatively assay sub-cellular acyl-CoA distribution. This approach builds on the SILEC approach using ^15^N_1_^13^C_3_-vitamin B5 (pantothenate) to generate cell lines highly enriched for ^15^N_1_^13^C_3_-labeled acyl-CoAs (Basu et al., 2011; Snyder et al., 2014). SILEC-SF applies SILEC-labeled cells as rigorous isotope labeled analogs used as internal standard controls introduced prior to fractionation to quantify metabolites in sub-cellular fractions. Thus, in addition to correcting for a range of factors in analysis by liquid chromatography-mass spectrometry (LC-MS) including variability in processing, analyte loss, extraction inefficiency, and ion suppression (Ciccimaro and Blair, 2010; Mann, 2006; Ong, 2012), SILEC-SF can also account for metabolic disruptions and sample losses during the fractionation process. Thus, SILEC-SF accounts for multiple confounding factors affecting sub-cellular fractionation, sample preparation and analysis. We applied SILEC-SF to the analysis of mitochondrial, cytosolic and nuclear metabolites and reveal distinct metabolic regulation of these compartments.

## Results

### SILEC-SF uses whole cell internal standards to control for sample processing

We hypothesized that inclusion of internal standards throughout fractionation and processing would enable accurate quantification of metabolites in sub-cellular compartments. SILEC with isotope labeled (^15^N_1_^13^C_3_-)vitamin B5 results in isotope label incorporation into the CoA backbone, which can be detected in acyl-CoA thioesters by LC-MS **(Figure 1A**)(Basu et al., 2011; Snyder et al., 2014). Taking advantage of the fact that this allows generation of ^15^N_1_^13^C_3_ labeled internal standards at >99% efficiency within living cells, we designed SILEC-subcellular fractionation (SILEC-SF) to achieve acyl-CoA quantitation in subcellular compartments **(Figure 1B).** For this, SILEC cells are harvested in fractionation buffer and then added in equal quantity to each experimental sample, which can be cells or tissues. After SILEC cell addition, cells are separated into subcellular compartments by fractionation. This strategy ensures the internal standard is present as whole cells before any processing for fractionation occurs, as well as in every relevant compartment after fractionation. In this way, the internal standards can account for sample loss that occurs from cell harvest to metabolite extraction. Each fraction is then extracted and analyzed separately by LC-MS and the ratio of the light (acyl-CoA molecules from experimental cells) to heavy (SILEC isotope labeled internal standard molecules) signal intensity is used to determine the quantities within each subcellular compartment across different experimental samples. Co-elution and simultaneous analysis of the analyte with the ^15^N_1_^13^C_3_ labeled internal standard improves quantitative performance (Frey et al., 2016). This approach is complementary to and can be adapted for use with a variety of fractionation approaches including differential centrifugation and isolation of organelles by immunoprecipitation **(Figure 2A & 2F).** Thus, SILEC-SF applies internal standards as living whole cells to enable rigorous metabolite quantification in subcellular compartments.

**Figure 1:**
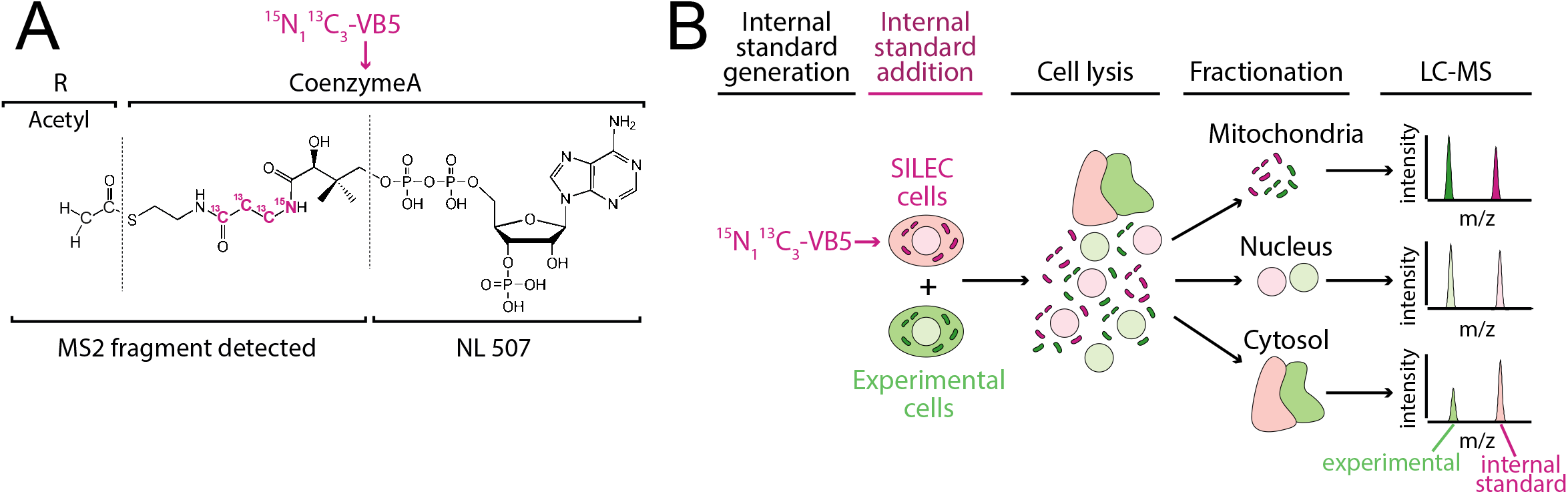
SILEC-SF uses whole cell internal standards to reveal compartment specific acyl-CoA profiles. **A)** ^15^N_1_^13^C_3_-isotope labeled vitamin B5 (VB5) is incorporated into coenzyme A (CoA) such that acyl-CoA species across all acyl (R group) species are isotope labeled. B) Representation of SILEC-SF workflow: internal standards were generated through isotope labeling, internal standard was added to samples as whole cells prior to cell lysis and separation of sub-cellular compartments by fractionation. The analyte and internal standard in each fraction was analyzed simultaneously by LC-MS and relative quantities are determined between samples.

**Figure 2:**
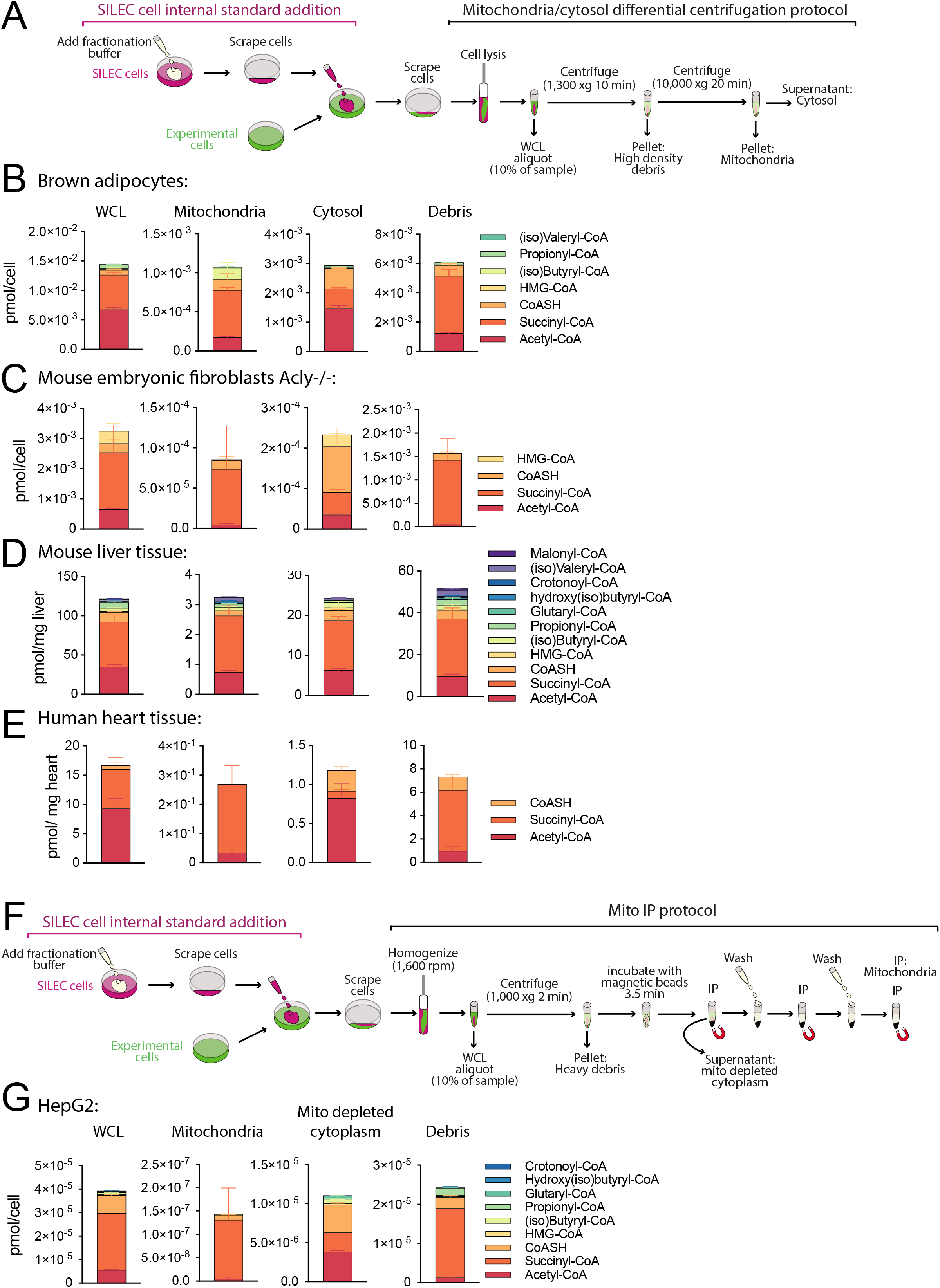
SILEC-SF reveals compartment-specific acyl-CoA profiles. A) Graphical representation of differential centrifugation method for mitochondria and cytosol isolation. B-E) Acyl-CoA quantitation in whole cell lysate (WCL), mitochondria, cytosol and high density debris (remainder material). B) Brown adipocytes in cell culture (n=4 replicate dishes) **C)** *Ady-/-* mouse embryonic fibroblasts (MEFs) were incubated in DMEM supplemented with 10% dialyzed FBS and 1 mM acetate for 4 h before cell harvest (n=4 replicate dishes) D) Mouse liver tissue (n=6 mice) D) Transmural left ventricle of human heart (n=5 replicate samples from a single heart). F) Graphical representation of Mito-IP protocol with SILEC-SF **G)** SILEC-SF using Mito-IP method for fractionation was applied to HepG2 cells incubated in serum free DMEM (5 mM glucose, 2 mM glutamine) for 2 h before harvest. Mean values from replicate samples (n=3) are displayed and error bars show standard deviation. Data for all acyl-CoA species that were quantified in each fraction are displayed. Those that were not quantified showed insufficient signal intensity for the analyte, the internal standard or both. Some metabolites indicated in the legend for C-G were not quantified in the mitochondrial fraction. These were B) HMG-CoA, **D)** Malonyl-CoA and **E)** CoASH, **G)** all except for acetyl-CoA succinyl-CoA and CoASH. (iso)Butyryl-CoA = Butyryl-CoA/lsobutyryl-CoA, (iso)Valeryl-CoA = Valeryl-CoA/lsovaleryl-CoA (isomers are not distinguished in analysis). HMG-CoA = 3-hydroxy-3-methylglutaryl-CoA.

### Standard curves specific to each fraction account for matrix-specific effects and allow absolute quantitation

We first established standard curves to achieve absolute quantitation of each metabolite recovered within each fraction. By using separate standard curves within each fraction, accurate quantitative comparisons can be made within each compartment across samples. Standard curves were generated by fractionation of additional aliquots of SILEC internal standard cells in parallel with experimental samples, followed by addition of known quantities of unlabeled standards **(Figure S1A & S1B).** Standard curves plot the light:heavy signal intensity ratio against quantity of unlabeled standard. Curves were strikingly different between subcellular fractions **(Figure S1C).** These distinct standard curves likely reflect several factors that vary across different fractions including the relative enrichment of specific acyl-CoA species (i.e. different quantities of SILEC internal standard), extraction efficiency, and matrix effects (Ciccimaro and Blair, 2010).

We next assessed the impact of matrix effects. Since the matrix is defined as all the components of the sample except the analyte (Guilbault and Hjelm, 1989), and those components are unique to each fraction, we reasoned that the effects of the matrix might be different across fractions. To examine this, we measured the impact of different sub-cellular matrices on raw signal intensity. SILEC sub-cellular matrices were generated by fractionation of fully labeled SILEC internal standard cells, similar to standard curve generation **(Figure S1D,** Western blots: **Figure S2B).** Unlabeled acyl-CoA standards, however, were added to the SILEC matrix extracts immediately before LC-MS analysis, as opposed to before extraction, to specifically assay the effects of matrix. Signal intensity for unlabeled acyl-CoA standards varied with the addition of different matrices in a manner that was specific to each metabolite **(Figure S1D).** This demonstrates that matrix-specific effects contribute to standard curve variation and highlight the importance of using matrix matched standard curves.

### Distinct acyl-CoA profiles define mitochondria and cytosol in diverse cell and tissue types

SILEC-SF was applied to determine the acyl-CoA distribution in mitochondria and cytosol in diverse cells and tissues including adipocytes in culture, fibroblasts, mouse liver, and human heart **(Figure 2B-E).** For maximal adaptability across sample types, we employed a classical differential centrifugation protocol for isolation of mitochondria and cytosol simultaneously (Clayton and Shadel, 2014; Frezza et al., 2007), with adaptations for speed and purity **(Figure 2A,** Western blots: **Figure S2B-E).** The cellular debris is enriched in high-density cellular material including unbroken cells and nuclei **(Figure 2A, Figure S2B-E).** SILEC-SF reveals distinctive acyl-CoA profiles defining cytosol and mitochondria. Succinyl-CoA, acetyl-CoA, and CoASH are the most abundant short chain acyl-CoA species on a whole cell level, consistent with data from direct extraction of matching whole cells and tissues in this study and previous studies **(Figure S2A)**(Bedi et al., 2016; Sadhukhan et al., 2016; Simithy et al., 2017). Notably, CoASH was enriched in the cytosol and succinyl-CoA was generally the dominant acyl-CoA species in the mitochondria, while acetyl-CoA was at similar or greater abundance than succinyl-CoA in the cytosol across these different cell/tissue types **(Figure 2B-E).** One of the strengths of SILEC-SF is that it can be applied in conjunction with different fractionation strategies. We therefore also adapted SILEC-SF to the recently developed mitochondrial immunoprecipitation method (Chen et al., 2017) **(Figure 2F & G),** for which we generated matrix specific standard curves by standard addition **(Figure S1B).** SILEC-SF was also successful in conjunction with this fractionation strategy, similarly identifying succinyl-CoA as prominent in the mitochondria and acetyl-CoA enriched outside of mitochondria **(Figure 2G).** Thus, the SILEC-SF method discerns distinct acyl-CoA profiles in mitochondria versus cytosol.

### SILEC-SF detects distinct mitochondrial adaptation to hypoxia

We next sought to validate the SILEC-SF method using a biologically relevant perturbation that would impact acyl-CoAs in sub-cellular compartments in a predictable manner. Reductive carboxylation of α-ketoglutarate (αKG) occurs in the mitochondria under hypoxia, whereby carbons from glutamine are directed to citrate production and away from succinyl-CoA production (Metallo et al., 2011) **(Figure 3A).** Thus, reduced mitochondrial succinyl-CoA levels were anticipated under hypoxia. SILEC-SF was applied to the analysis of succinyl-CoA in the mitochondria and cytosol in HepG2 cells incubated under 20% (normoxia) and 1% (hypoxia) oxygen for 24 hours. We first assessed the effect of hypoxia exposure on acyl-CoA levels in whole HepG2 cells with rapid direct extraction. Succinyl-CoA was substantially reduced (~50%) under hypoxia, whilst acetyl-CoA was minimally impacted **(Figure 3B).** SILEC-SF analysis revealed that succinyl-CoA was reduced specifically in the mitochondrial compartment in hypoxia **(Figure 3C).** In contrast, cytosolic succinyl-CoA levels were unchanged by hypoxia, indicating an independently regulated metabolic pool **(Figure 3C).** Mitochondrial acetyl-CoA was also markedly reduced by hypoxia despite a lack of change in whole cell pools, consistent with the majority of cellular acetyl-CoA being cytosolic and demonstrating the importance of sub-cellular analysis to uncover metabolic changes in compartments that contain smaller pools of a given metabolite. We also assessed compartmentalized abundance of a panel of short chain acyl-CoA species **(Figure 3D,** representative data in pmol/cell: **Figure S3A),** finding that hypoxia resulted in reduced abundance of multiple acyl-CoAs in mitochondria but not in cytosol. These data demonstrate that SILEC-SF detects compartment-specific changes in acyl-CoA abundance in response to biological perturbation.

**Figure 3:**
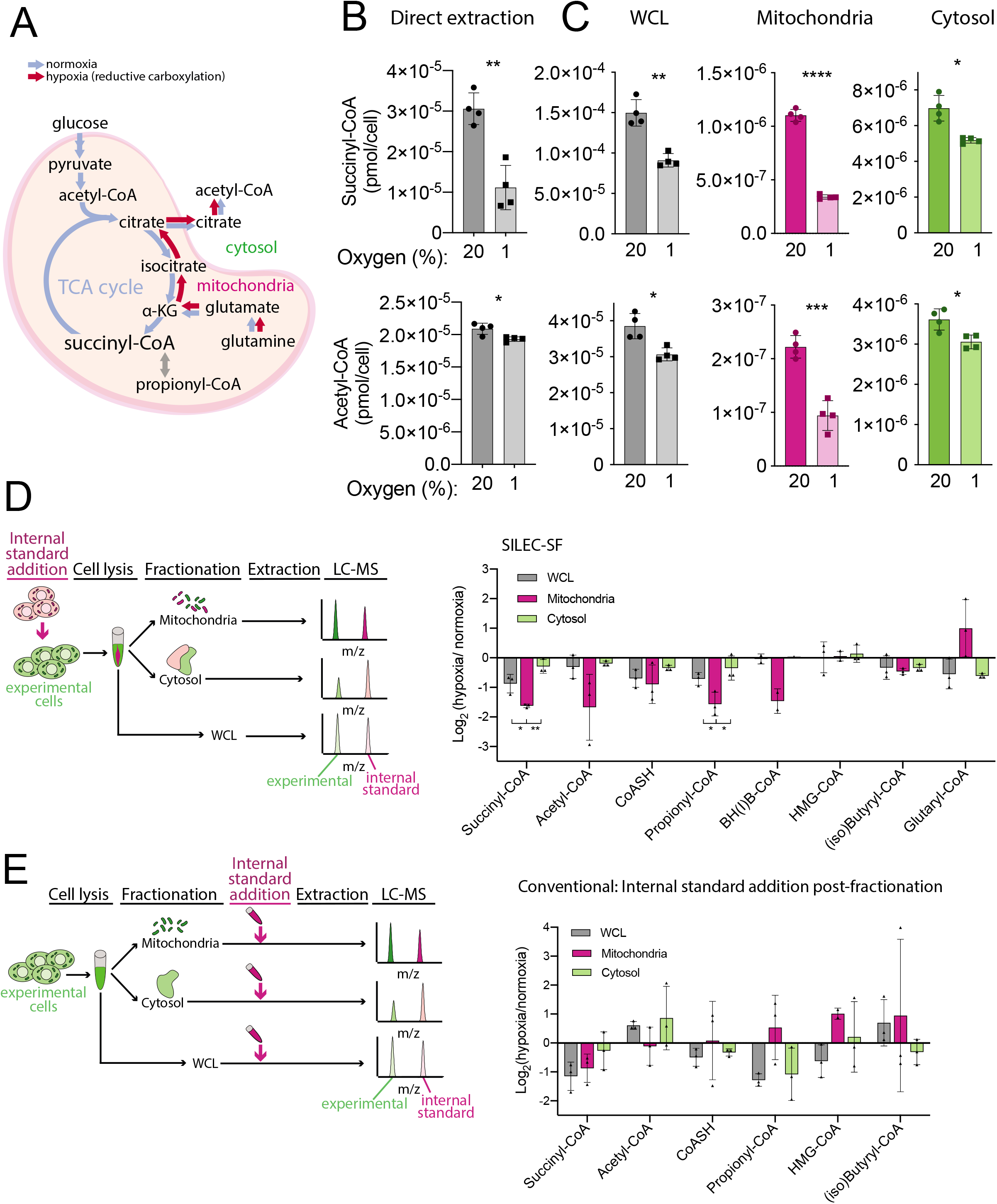
SILEC-SF detects distinct mitochondrial adaptation to hypoxia. **A)** Schematic comparing glutamine metabolism in the TCA cycle under hypoxic and normoxic conditions. B-E) HepG2 cells were incubated under 20% (normoxia) or 1% (hypoxia) for 24 h. Cells were serum starved in DMEM containing 5 mM glucose and 2 mM glutamine under their respective oxygen tensions for 2 h before harvest. Serum starvation was used to avoid potential variability in metabolic response to serum batches, which could vary between experiments. Symbols indicate individual replicate dishes (n=4) from a representative experiment. B) Whole cell analysis after direct extraction of metabolites. **C)** Succinyl-CoA and acetyl-CoA quantitation by SILEC-SF using the mitochondrial/cytosolic differential centrifugation procedure from a representative experiment (a replicate of that displayed in **Fig 5C,** that is also incorporated into **Fig 3D** and **Fig 5E). D,E)** Fold-change (hypoxia/normoxia) from mean absolute quantitation as pmol/cell were calculated for 3 independent experiments conducted on separate days and log transformed. Symbols indicate the mean from each experiment. D) SILEC-SF involves introduction of internal standard before fractionation. E) Schematic for internal standard addition after fractionation. Abbreviations: BH(l)B-CoA =3-Hydroxybutyryl-CoA/ 3-Hydroxyisobutyryl-CoA, (iso)Butyryl-CoA = Butyryl-CoA/lsobutyryl-CoA (isomers are not distinguished in analysis). For all graphs, error bars represent standard deviation. Statistical comparisons were made by two-tailed Student’s ř-test with Welch’s correction and statistical significance was defined as p < 0.05 (*), p < 0.01 (**), p < 0.001 (***), p < 0.0001 (****).

### SILEC-SF improves on internal standard addition after fractionation

To further validate the method, we next tested the extent to which SILEC-SF: 1) improves on metabolite quantification over conventional addition of internal standards after fractionation, and 2) accounts for sample loss and metabolic activity during processing. For these tests, we leveraged the strong compartmentalized response to hypoxia. First, to assess the effectiveness of SILEC-SF compared to introduction of conventional addition of internal standard after fractionation **(Figure 3D & 3E),** data for each method were compared across 3 separate experiments carried out on separate days. Although the conventional method reproduced the core result of mitochondrial-specific succinyl-CoA reduction under hypoxia, the variability was greater and data on other lower abundance acyl-CoAs was inconsistent between experiments. Therefore, SILEC-SF improves quantification of compartmentalized acyl-CoA pools over conventional addition of internal standard after fractionation.

We next asked whether SILEC-SF accounts for metabolic activity during processing. We previously demonstrated that post-harvest metabolism occurs during fractionation by assessing the incorporation of isotope labeled substrates introduced in the fractionation buffer into downstream metabolic products (Trefely et al 2019). We applied post-harvest labeling to assess the extent to which SILEC internal standards account for post-harvest metabolism **(Figure S3B).** ^13^C_5_-glutamine was used since it is an important substrate for succinyl-CoA generation. ^13^C_5_-glutamine was incorporated as an additional ^13^C_4_-label into succinyl-CoA M4 (unlabeled CoA derived from experimental cells) and into succinyl-CoA ISTD M4 (heavy CoA derived from internal standard cells). Differences in post-harvest metabolism were apparent between fractions (5-11% in mitochondria and debris versus <1% in cytosol), but were comparable within each fraction in the experimental and internal standard cells. This indicates that SILEC internal standards undergo compartment-specific post-harvest metabolic changes mirroring those in experimental cells, and can thus account for these changes by retaining the differences between internal standard and experimental analyte across samples after spiking in the internal standard. Together these data indicate that SILEC-SF improves sub-cellular acyl-CoA quantitation, in part by accounting for post-harvest metabolism.

### Cytosolic HMG-CoA is exquisitely sensitive to acetate supply

Having validated that acyl-CoA abundance changes within mitochondria are reported by SILEC-SF, we next aimed to validate SILEC-SF for detection of changes in cytosolic acyl-CoA pools. For this, we leveraged two models of ATP-citrate lyase (ACLY) deficiency, which rely on acetate to supply cytosolic acetyl-CoA via Acyl-CoA Synthetase Short Chain Family Member 2 (ACSS2): *Acly^-/-^* MEFs (Zhao et al., 2016) and *Acly^-/-^* murine liver cancer cells **(Figure S4A).** Within the cytosol, acetyl-CoA is used for generation of malonyl-CoA for fatty acid synthesis and HMG-CoA in the mevalonate pathway for synthesis of sterols and isoprenoids **(Figure S4A).** We first investigated the relationship between acetate dose and acyl-CoA abundance in whole cells using direct rapid extraction of metabolites. Cells were incubated in 0, 0.1, or 1 mM acetate for 4 hours, and the dose response was compared across 8 short chain acyl-CoA species quantified **(Figure 4B, Figure S4B-C).** This analysis revealed that HMG-CoA is severely depleted upon acetate withdrawal and increases 10-40 fold with increasing acetate supplementation in *Acly^-/-^* cells **(Figure 4C).** Acetyl-CoA was also sensitive to exogenous acetate, but the concentration changed by only 2-3 fold in the presence or absence of acetate **(Figure 4C).** In contrast, malonyl-CoA did not exhibit clear responsivity to acetate in whole cells **(Figure 4B, Figure S4B-D).** Acyl-CoA levels in control (*Acly^f/f^*) cells are largely unaffected by acetate supplementation **(Figure 4B-C, Figure S4D).**

**Figure 4:**
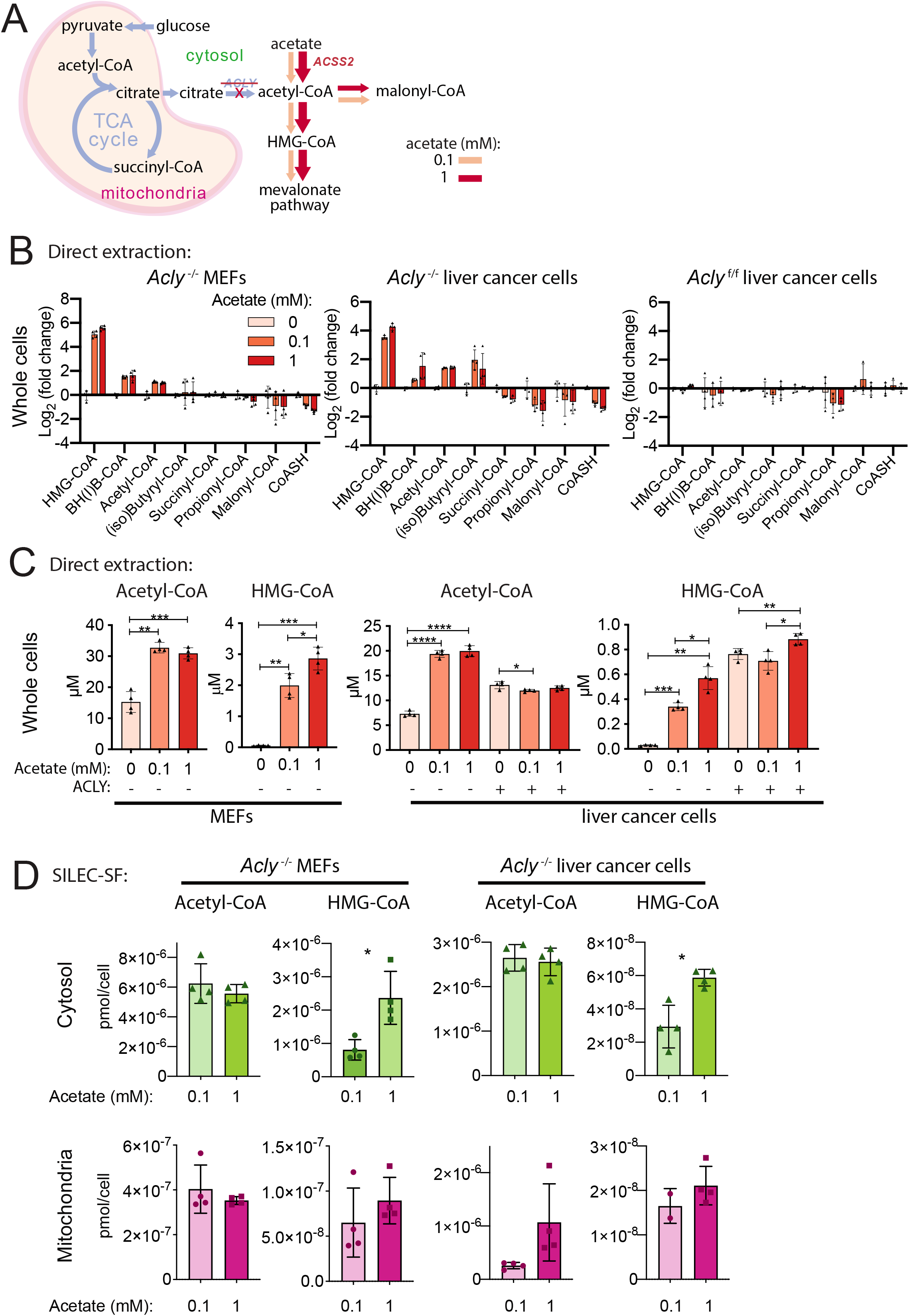
Cytosolic HMG-CoA is a sensitive readout of cytosolic acetate supply. **A)** Schematic representation of cytosolic acetyl-CoA generation. Acetate supplies cytosolic acetyl-CoA through upregulation of ACSS2 in ACLY deficient cells. B-D) Cells were incubated for 4 h in DMEM supplemented with 10% dialyzed fetal calf serum with the addition of the indicated amount of acetate for 4 h. Whole cell acyl-CoA concentrations were determined in ACLY deficient mouse embryonic fibroblasts *(Acly^-/-^* MEFs) and liver cancer cell line *(Acly^-/-^* liver cancer cells) as well as matched ACLY sufficient control *(Acly^f/f^* liver cancer). Symbols represents individual replicate cell dishes (n=4) from representative experiments. B) Whole cell direct extraction fold-change analysis. Fold change compared to mean 0 mM acetate concentration (in μM) was calculated and log transformed. C) Whole cell direct extraction (data from B) displayed as cellular concentration for acetyl-CoA and HMG-CoA. D) SILEC-SF was used to specifically determine cytosolic acyl-CoA response to 0.1 versus 1 mM acetate supplementation. For all panels, error bars show standard deviation and statistical comparisons between two groups, were made by two-tailed Student’s *t*-test with Welch’s correction and statistical significance defined as p < 0.05 (*), p < 0.01 (**), p < 0.001 (***), p < 0.0001 (****).

**Figure 5:**
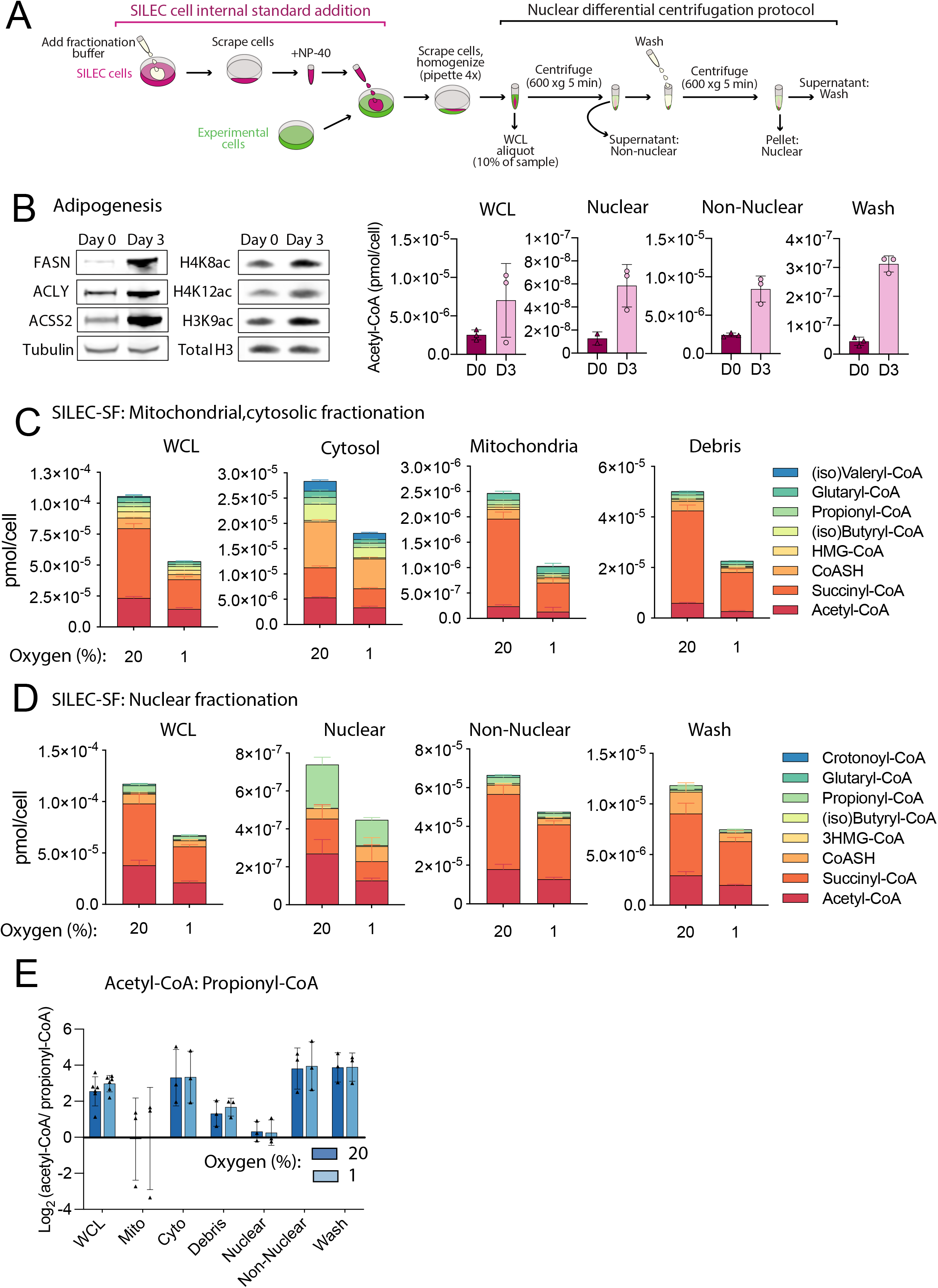
SILEC-SF identifies enrichment of propionyl-CoA in the nucleus. **A)** Graphical representation of differential centrifugation method for nuclear isolation. B) Preadipocytes (5A) were harvested at day 0 and day 3 following induction with differentiation cocktail. Western blots of acid extracted histones and non-nuclear fraction. Representative experiment comparing profiles of short chain acyl-CoA species quantified in each fraction. C,D) HepG2 cells were incubated under 20% (normoxia) or 1% (hypoxia) for 24 h. Cells were serum starved in DMEM containing 5 mM glucose and 2 mM glutamine under their respective oxygen tensions for 2 h before harvest to maintain stable conditions across different experiments. **C)** Profiles of short chain acyl-CoA species quantified after SILEC-SF using the mitochondrial/cytosolic differential centrifugation procedure from a representative experiment (a replicate of that displayed in **Figure 3C,** that is also incorporated into **Figure 3D** and **Figure 5E)**. Some metabolites indicated in the legend for **C** were not quantified in all fractions, specifically, crotonoyl-CoA was quantified only in non-nuclear fraction, and glutaryl-CoA, (iso)butyryl-CoA, and HMG-CoA were not quantified in the nuclear fraction. Those that were not quantified showed insufficient signal intensity for the analyte, the internal standard or both. D) Profiles of short chain acyl-CoA species quantified after SILEC-SF using the nuclear differential centrifugation procedure from a representative experiment (a replicate which is also incorporated into **Figure 5E).** n=4 replicate cell dishes/experiment. E) Fold-change (propionyl-CoA/acetyl-CoA) was calculated from absolute quantitation (pmol/cell) for 3 independent experiments conducted on separate days (n=4 individual replicate cell dishes within each experiment). Symbols indicate mean from each experiment. For all graphs, error bars show standard deviation.

We predicted that the dose-responsive change in HMG-CoA abundance occurs specifically in the cytosol, since our prior sub-cellular tracing analyses demonstrated a close kinetic relationship between cytosolic acetyl-CoA and HMG-CoA labeling from acetate (Trefely et al., 2019). We performed SILEC-SF to compare acyl-CoA levels specifically in the cytosolic and mitochondrial compartments in response to exogenous acetate **(Figure 4D).** Importantly, the direct extraction and SILEC-SF whole cell data reported consistent trends **(Figure S4B,C,E,F).** Acetyl-CoA levels were not significantly different between low and high acetate in either mitochondrial or cytosolic compartments **(Figure 4D).** HMG-CoA, however, responded to acetate dose specifically in the cytosolic and not in the mitochondrial compartment of *Acly^-/-^* cells **(Figure 4D).** The significant response of cytosolic HMG-CoA to acetate dose was unique amongst a panel of 6 to 7 short chain acyl-CoA species quantified in the cytosol of *Acly^-/-^* MEF and liver cancer cells **(Figure S4E & S4F).** Together, the data indicate that SILEC-SF can report on perturbations to cytosolic acyl-CoA pools and indicate that in the absence of ACLY, cytosolic HMG-CoA is highly sensitive to and dependent on exogenous acetate availability.

### SILEC-SF reveals distinct nuclear acyl-CoA profiles, including enrichment of propionyl-CoA

We next applied SILEC-SF to the quantification of acyl-CoAs within the nucleus, with the goals of understanding: 1) if the nucleus is distinct metabolically from the cytosol and 2) if this approach can be used to gain novel insights into nuclear metabolism and its links to the chromatin modification. Nuclear metabolite quantitation is a particular challenge due in part to large nuclear pores through which small molecules can diffuse. We reasoned that this leak could be accounted for by parallel changes in internal standard nuclei in the SILEC-SF approach.

We applied SILEC-SF to nuclear acyl-CoA determination, using a rapid differential centrifugation protocol to separate nuclei from non-nuclear fractions **(Figure 5A).** We first applied this to quantitation of nuclear-specific acetyl-CoA in an adipocyte differentiation model, since histone acetylation levels and whole cell acetyl-CoA abundance increase markedly during this process (Fernandez et al., 2019; Wellen et al., 2009) **(Figure 5B).** A 5-fold increase in nuclear acetyl-CoA was measured upon differentiation induction (day 3 versus day 0), correlating with histone acetylation. Perhaps unsurprisingly, acetyl-CoA levels increased during differentiation not only in the nucleus, but also in whole cell lysate and the non-nuclear fraction **(Figure 5B).** Thus, the adipocyte differentiation model documented a correlation between nuclear acetyl-CoA abundance and histone acetylation, but did not allow for discernment of whether nuclear acyl-CoA profiles are distinct from that in other compartments.

To further investigate the potential for nuclear-specific acyl-CoA regulation, we applied SILEC-SF to nuclear acyl-CoA quantification in the HepG2 hypoxia model **(Figure 5D, Figure S5A-D;** Western blots: **Figure S3C).** To compare nuclear acyl-CoA pools to those in other compartments, we first confirmed that whole cell lysate data from the nuclear and mitochondrial fractionation protocols are consistent, indicating the validity of comparisons across these experiments **(Figure 5C & 5D).** Remarkably, comparison of acyl-CoA quantification across different fractions revealed that the nuclear metabolite profile was distinct from whole cells and other compartments including the cytosol **(Figure 5C & 5D)** Notably, propionyl-CoA is substantially enriched in the nucleus compared to the cytosol. This is emphasized by examining acetyl-CoA: propionyl-CoA ratio across sub-cellular fractions in which data from the mitochondria/cytosol and nuclear/non-nuclear fractionation methods were combined from 3 independent experiments carried out on different days for each method **(Figure 5E).** Acetyl-CoA and propionyl-CoA are approximately equimolar in the nucleus, whereas in the whole cell lysate, cytosolic and non-nuclear fractions, acetyl-CoA is more abundant. The non-nuclear acyl-CoA profile closely reflected that of the whole cell lysate, indicating that the nuclear acyl-CoA pool did not contribute substantially to the whole cell acyl-CoA abundance **(Figure 5D, Figure S5A & S5B).** The wash fraction also reflected the non-nuclear and whole cell lysate **(Figure 5D, Figure S5A & S5D),** suggesting that the wash was necessary to remove residual non-nuclear material from the purified nuclear pellet. Additionally, the profile of acyl-CoAs in the mitochondrial fraction and the nuclear fraction were very distinct **(Figure 5C** vs **Figure 5D),** highlighting that mitochondrial contamination was not likely to explain the nuclear acyl-CoA pool. Thus, SILEC-SF reveals distinct and reproducible differences in nuclear acyl-CoA profiles versus other compartments. Furthermore, these data identify a unique enrichment for propionyl-CoA within the nucleus.

### Isoleucine catabolism contributes to nuclear propionyl-CoA generation and histone lysine propionylation (Kpr) marks

We next addressed the metabolic source of nuclear propionyl-CoA. Pathways for endogenous propionyl-CoA generation are annotated to the mitochondria, where its metabolic fate is to enter the TCA cycle via succinyl-CoA **(Figure 6A),** although it may also leave the mitochondria via the carnitine shuttle (Trefely et al., 2020). To investigate the metabolic origin of nuclear propionyl-CoA, we first assessed the substrate contribution to total cellular propionyl-CoA by stable isotope tracing of a panel of uniformly ^13^C-labeled substrates followed by direct extraction of whole cells. Strikingly, isoleucine labeling into propionyl-CoA M3 contributed 30%, 43% and 50% of the propionyl-CoA pool, in HepG2, HeLa and murine pancreatic ductal adenocarcinoma cells (KPC cells), respectively, in the absence of propionate **(Figure 6B).** This accounts for the majority of cellular propionyl-CoA since the tracer was 50% diluted with unlabeled isoleucine. In contrast, the contribution of isoleucine to the cellular acetyl-CoA pools was <1%, and to succinyl-CoA pools varied across cell lines (<1% to 15%) **(Figure S6A).** Propionate also fed into propionyl-CoA pools in all 3 cell lines when supplemented as a positive control **(Figure 6B).** Since isoleucine is the dominant substrate for propionyl-CoA generation under these conditions, we tested the potential for isoleucine to contribute specifically to nuclear propionyl-CoA using sub-cellular kinetic analysis with isotope postlabeling to correct for post-harvest metabolic activity (as described by (Trefely et al., 2019)). Nuclear propionyl-CoA was labeled ~20% by uniformly labeled ^13^C_6_-isoleucine 50% diluted with unlabeled isoleucine in KPC cells, indicating that the propionyl-CoA pool in the nucleus can be substantially derived from isoleucine **(Figure 6C).** Thus, nuclear propionyl-CoA pools are derived, at least in part, from BCAA catabolism.

**Figure 6:**
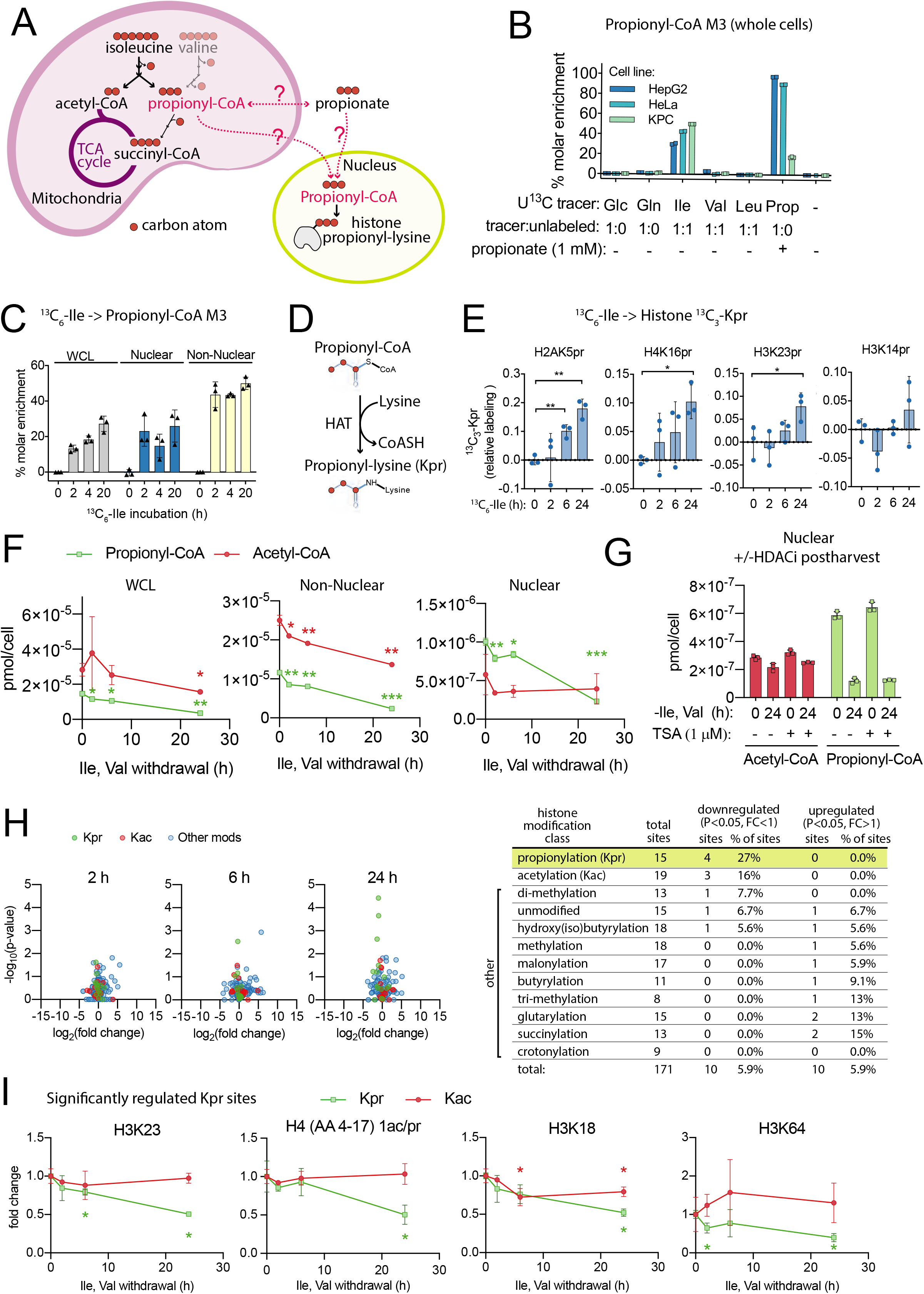
Isoleucine catabolism is a major substrate for nuclear propionyl-CoA generation and histone lysine propionylation. **A)** Schematic representation of propionyl-CoA generation. Propionyl-CoA is generated in the mitochondria from multiple sources including isoleucine and valine. 3 of the 6 carbons in lie contribute to the acyl group of propionyl-CoA. B) Incorporation of various substrates into propionyl-CoA, was compared in whole cells by direct extraction of HepG2, HeLa and pancreatic adenocarcinoma (KPC) cells incubated in media containing uniformly (U) ^13^C-labeled substrates for 18 h. The total concentration of each substrate was equal across all samples except for propionate, which was added only to the U^13^C_3_-propionate samples. U^13^C-labeled Val, Leu and lie were diluted 1:1 with unlabeled substrate. N=3 replicate samples per condition from a single experiment. Error bars are standard deviation. **C)** KPC cells were incubated in media containing uniformly labeled ^13^C_6_-isoleucine diluted 1:1 with unlabeled isoleucine. Fractionation was performed with post-labeling in the presence of partially labeled ^13^C_6_-isoleucine to account for post-harvest metabolism. Incorporation of ^13^C_6_-isoleucine was determined by enrichment of propionyl-CoA M3 at the indicated incubation times. D) Schematic representation of histone lysine propionylation catalyzed by histone acyl transferase (HAT) enzymes. E) HepG2 cells were incubated in DMEM supplemented with 10% dialyzed FBS (to avoid propionate contamination) in which isoleucine was entirely replaced with ^13^C_6_-isoleucine for the indicated times. Histone acyl proteomic analysis was performed on acid extracted histones. 4 distinct Kpr containing peptides were identified with sufficient signal intensity for both labeled (^13^C_3_.Kpr) and unlabeled Kpr and data are represented as relative labeling normalized to unlabeled controls (t=O). F-l) HepG2 cells were incubated in DMEM with 10% dialyzed FBS. Media was changed on all timepoint at t=-24 so that they all experienced the same formulation, and switched to the lie and valine dropout media at the indicated times before harvest. F) SILEC-SF analysis with nuclear/non-nuclear fractionation. G) Samples were treated with 1 μM trichostatin A (TSA) in fractionation buffer. H,l) proteomic analysis of acid extracted histones. H) volcano plots compare intensity of specific histone peptide modifications for each time point to t=0 control. Table summarizes histone modification classes and sites that were significantly regulated at any timepoint. Single and double modifications of a peptide that could not be assigned to a specific residue are counted as distinct sites. I) relative intensity over time for 4 Kpr sites identified as significantly regulated at any timepoint and for corresponding Kac marks. Statistical significance in E & F was determined by comparison to t=O control for each mark or metabolite and is indicated above or below the specific timepoint in the color corresponding to the data set. p < 0.05 (*), p < 0.01 (**), p < 0.001 (***).

Since acyl-CoAs are substrates for histone lysine acylation marks such as acetylation and propionylation, we hypothesized that a fate of isoleucine derived propionyl-CoA in the nucleus could be as a substrate for histone propionylation **(Figure 6D).** To directly interrogate this, we examined the incorporation of U^13^C_6_-isoleucine into histone propionyl-lysine (Kpr) marks by mass spectrometry. ^13^C_3_ label incorporation into Kpr was measured using the 4 best detectable peptides modified with endogenous propionylation (H2AK5pr, H4K16pr, H3K23pr and H2K14pr). The signal extraction was not feasible for other potentially propionylated peptides, given the low abundance of the modification and the even lower abundance of peptide isotopes. Increasing label incorporation was observed in H2AK5pr, H4K16pr and H3K23pr, but not in H2K14pr, in cells incubated over a 24 h time-course in ^13^C_6_-isoleucine **(Figure 6E),** demonstrating the direct contribution of propionate acyl-chains with all three carbons from isoleucine into histone Kpr marks.

### Nuclear propionyl-CoA responds to isoleucine and valine supply

We hypothesized that BCAA availability would impact both nuclear propionyl-CoA and histone Kpr marks since isoleucine directly supplies carbons for both. To test this, cells were incubated over a time course in media lacking isoleucine and valine. Nuclear acyl-CoA analysis by SILEC-SF revealed a substantial (4-5 fold) decrease in propionyl-CoA after 24 h of deprivation of these amino acids **(Figure 6F & 6G).** Propionyl-CoA levels in the whole cell and non-nuclear fraction mirrored the nuclear trends **(Figure S6B).** Nuclear acetyl-CoA levels, in contrast, were only modestly impacted under these conditions **(Figure 6F & 6G).** These data indicate nuclear propionyl-CoA pools are sensitive to the availability of BCAAs.

One possible artifactual explanation for the detection of high nuclear propionyl-CoA abundance is that it results from depropionylation through the action of histone deacetylases (HDACs)(McClure et al., 2017) during processing. The reaction catalyzed by class I and II HDACs converts lysine-acyl marks to their corresponding fatty acid, which could potentially be acted upon by nuclear resident acyl-CoA synthetase enzymes to produce the corresponding acyl-CoA (Bulusu et al., 2017). Deacylation by class III HDACs (sirtuins) produces O-acyl-ADP-ribose, which also has the potential to be converted to fatty acids and acyl-CoAs, although more enzymatic steps are required (Chen et al., 2011; Tong and Denu, 2010). To test potential for this mechanism to generate propionyl-CoA or acetyl-CoA during SILEC-SF the class I and II HDAC inhibitor, trichostatin A (Bradner et al., 2010; Khan et al., 2008), was added to the fractionation buffer and compared to a side-by side control experiment **(Figure 6G, Figure S6B).** Our results indicate no contribution from HDAC dependent deacylation to acyl-CoA detection during fractionation, corroborated by Western blots **(Figure S5C).** The data indicate that artifactual deacylation during processing does not contribute detectably to nuclear acyl-CoA profiles, supporting the conclusion that nuclear acyl-CoA profiles are biologically distinct from those in other compartments.

### Isoleucine and valine supply controls selected histone lysine propionylation (Kpr) marks

Since nuclear propionyl-CoA levels were responsive to isoleucine and valine withdrawal, we used this as a model to test the functional relevance of nuclear propionyl-CoA in regulating histone Kpr. Western blot analysis of acid extracted histones using pan Kpr and lysine acetylation (Kac) antibodies showed no notable change in either modification **(Figure S6D).** This was surprising given the substantial metabolic shift detected in nuclear propionyl-CoA and raised the possibility that there might be cross reactivity in the Kpr antibodies (Simithy et al., 2017) and/or selective sensitivity in Kpr marks to isoleucine availability.

Acyl proteomic analysis (Sidoli et al., 2019a) was employed to analyze specific histone peptide modifications. A total of 171 distinctly modified histone peptides comprising 10 different chemical modifications across 15 histone peptides were quantified, including 15 distinct histone Kpr sites **(Figure 6H, Figure S6E).** Kpr marks were proportionally the most significantly regulated of all marks with 27% of quantified Kpr sites being significantly downregulated. No Kpr sites were significantly upregulated. 4 distinct histone Kpr sites were identified as significantly downregulated (H3K23, H3K18, H3K64 and the H4 peptide from amino acid 4 to 17). This histone H4 peptide contains four possible modifiable residues (K5, K8, K12 or K16). This analysis did not discriminate which residue was propionylated, as it would require quantifying isotopes at the MS/MS product ion level, for which we do not have sufficient sensitivity in this analysis **(Figure 6I).** The progressive loss of Kpr at these sites over a 24 h timecourse correlates with nuclear propionyl-CoA levels **(Figure 6F).** Consistent with the relatively unchanged acetyl-CoA levels, the degree of change in Kac is small compared to Kpr. This is highlighted by comparing the Kac and Kpr regulation in the 3 sites with significantly downregulated Kac **(Figure 6I, Figure S6F).** Ranking of Kpr marks by average % intensity gives a semi-quantitative indication of their abundance **(Figure S6G).** The Kpr sites top ranked by signal intensity are not regulated by isoleucine and valine withdrawal, consistent with the pan Kpr antibody data and highlighting the importance of site-specific analyses to uncover metabolic regulation. Thus, nuclear propionyl-CoA supply is sensitive to the supply of BCAAs and regulates Kpr abundance at select histone lysine sites.

## Discussion

In this study we present SILEC-SF, a rigorous isotope dilution approach to sub-cellular quantitation of acyl-CoAs. This represents a critical advance that enables direct analysis of the mechanistic relationship between acyl-CoA supply and functional outputs of acyl-CoA metabolism including protein acylation and metabolic pathway activity, which are distinct between sub-cellular compartments.

Quantification of compartmentalized metabolite pools has been previously approached by several methods. Genetically encoded fluorescent metabolite sensors can be targeted to specific compartments, and have been applied to a limited set of metabolites, including nicotinamide adenine dinucleotides (NAD+, NADH), adenosine triphosphate (ATP), glucose, glutamine, lactate, pyruvate, S-adenosyl methionine (SAM) and guanosine triphosphate (GTP) (Jaffrey, 2018; Okumoto et al., 2012; Zhang et al., 2018). However, these methods have technical limitations in that they require highly engineered experimental settings, and probes generally do not allow simultaneous monitoring of multiple metabolite species. Mass spectrometry (MS) has the advantage of direct and highly multiplexed analyses. Imaging MS holds great potential and has shown utility in measurement of intracellular drug concentration(Lee et al., 2017) as well as endogenous metabolites including dopamine in extracellular vesicles (Legin et al., 2014; Thomen et al., 2020), and membrane-specific measurements of cholesterol and lipid species including phosphor and glycolipids (Dueñas et al., 2017; Niehaus et al., 2019) but the processing, sensitivity and spatial resolution required to achieve robust organelle-specific quantitation for many metabolites is limiting (Bowman et al., 2020). Cellular fractionation can be scaled for appropriate quantitation of a range of biomolecules. However, sub-cellular analyses have been a major technical challenge in metabolomics due to requirements for sample processing and metabolite extraction. We have previously shown that post-harvest metabolism occurs during cell harvest and fractionation and can introduce artifacts that impact data interpretation (Trefely et al., 2019). The innovation of SILEC-SF mitigates these problems by introducing internal standards as whole cells before fractionation.

SILEC-SF is a flexible approach applicable to a variety of sub-cellular fractionation platforms and biological samples, and adaptable across multiple classes of metabolites. In this study we applied SILEC-SF to differential centrifugation to achieve quantification of metabolites in mitochondria, cytosol, and nucleus, and to immunoprecipitation of tagged mitochondria. Organelle immunoprecipitation approaches have been developed for lysosomes and peroxisomes, in addition to mitochondria (Bayraktar et al., 2019; Chen et al., 2016, 2017; Ray et al., 2020; Wyant et al., 2017; Xiong et al., 2019), opening the door for SILEC-SF to be applied to these organelles in the future. SILEC-SF may, in principle, be applied across additional platforms such as non-aqueous fractionation, generally applied to plant cells (Dietz, 2017; Fly et al., 2015; Krueger et al., 2014). SILEC-SF can be used to analyze a broad range of biological samples including human tissues for clinical studies, as demonstrated in this study,. Furthermore, SILEC is not limited to acyl-CoAs. The SILEC principle can be extended to other classes of metabolites that can be completely labeled in cell types of interest. For example, we have used SILEC labeling in mammalian cells to generate nicotinamide adenine dinucleotide (NAD+/NADH) and nicotinamide adenine dinucleotide phosphate (NADP/NADPH) labeled internal standards (Frederick et al., 2017). Thus SILEC-SF is a principle for rigorous internal standard controls applicable to sub-cellular metabolite quantitation by MS across multiple fractionation platforms and biological systems.

By applying SILEC-SF to nuclear analysis, we make the intriguing observation that propionyl-CoA is enriched within the nucleus relative to other compartments. This is compelling because histone lysine propionylation, which uses propionyl-CoA as a substrate, has been identified as a dynamically regulated chromatin modification associated with active gene transcription (Kebede et al., 2017; Lagerwaard et al., 2021; Liu et al., 2009; Simithy et al., 2017), although levels and sources of propionyl-CoA in the nucleus are unclear (Trefely et al., 2020). We identify isoleucine as a source of nuclear propionyl-CoA that also feeds into histone lysine propionylation. These findings open up numerous questions for future investigation. Firstly, what are the mechanisms of nuclear propionyl-CoA production? Isoleucine catabolism occurs in the mitochondria and mechanisms for transport of metabolic intermediates in this pathway to the nucleus are unknown. Carnitine acetyltransferase (CrAT) and the vitamin B12 dependent enzyme methylmalonyl-CoA mutase have been implicated in the supply of isoleucine derived propionyl-CoA for cytosolic fatty acid synthesis (Crown et al., 2015; Green et al., 2015; Wallace et al., 2018), and similar mechanisms could plausibly feed into nuclear propionyl-CoA pools. Secondly, how is nuclear enrichment of propionyl-CoA maintained versus the cytosol? Is propionyl-CoA specifically sequestered or generated within the nucleus, or are other acyl-CoA species preferentially consumed? In future studies, sub-cellular quantitation coupled with genetic manipulation of putative transport mechanisms will be useful to elucidate the spatial organization of metabolic pathways connecting isoleucine to nuclear propionyl-CoA.

Our discovery that Kpr marks at specific histone sites are correlated to nuclear propionyl-CoA levels, raises the question of how such specificity is achieved. One possibility is that enzymes responsible for propionyl-CoA generation are physically linked to multiprotein complexes targeted to specific histone sites as has been reported for histone succinylation (Wang et al., 2017). Another possibility is that acyl transferase enzymes sensitive to propionyl-CoA concentration (or relative concentration in competition with other acyl-CoA species) preferentially target specific histone sites. Another question is how these chromatin modifications are linked to cell function and disease, particularly since BCAAs play an important role in a multiple disease processes including cancer and insulin resistance (Neinast et al., 2019; Sivanand and Vander Heiden, 2020; White et al., 2021). Interestingly, recent evidence indicates that isoleucine is a major driver in the adverse metabolic response to dietary BCAAs (Yu et al., 2021). Future investigations will illuminate the roles isoleucine metabolism in regulating Kpr marks in disease processes.

In summary, SILEC-SF of provides a multiplexed approach that can be used to quantitatively measure sub-cellular metabolite pools. It allows the differentiation of environmental and nutrient responses between the whole cell, mitochondria, cytoplasm, and surprisingly the nucleus. Combined with orthogonal approaches of isotope tracing and proteomics, our data indicates that a nutrient specific response is elicited in the nucleus via propionyl-CoA with resulting changes in histone PTMs.

## Methods

### Cell Culture

Cells were maintained at 37 °C and 5% CO_2_ and passaged every 2-3 days at 80% confluence. All cells were tested and mycoplasma-free. Mouse embryonic fibroblast (MEF) cell lines were cultured in DMEM, high glucose (Thermo Fisher Scientific, Gibco #11965084) with 10% calf serum (Gemini bio-products #100-510, lot C93GOOH). Hepatocellular carcinoma (HepG2) were used at <20 passages and were cultured in DMEM, high glucose with 10% fetal bovine serum (Gemini Biosciences). Murine pancreatic adenocarcinoma cells were cultured in DMEM, high glucose (Thermo Fisher Scientific, Gibco #11965084) with 10% calf serum (Gemini bio-products #100-510, lot C93GOOH). Hepa1c7 cells were cultured in alpha MEM (Thermo Fisher Scientific, Gibco cat. #12561056) with 10% fetal bovine serum (Gemini Biosciences) and penicillin/streptomycin (Thermo Fisher Scientific, Gibco cat. #10378016). Brown adipocytes were prepared from immortalized brown preadipocyte cells (Harms et al., 2014) through induction of differentiation as described previously (Huber et al., 2019). 5A preadipocytes were passaged in DMEM/F12 (Gibco #11320033), 10% heat inactivated FBS, 1% Pen/Strep and seeded in 12 well plates for experiments. Cells were cultured for 24-48 h post confluent before induction media was added for 3 days. Induction media; DMEM/F12,10% heat inactivated FBS, 1% Pen/Strep, 5ug/ml insulin, 0.05mM IBMX, 10μM Dexamethasone, 5μM Troglitazone.

### Liver cancer cell line (Acly^f/f^ clone D42 and *Acly^-/-^* clone D42C4) generation

Acly^f/f^ liver cancer cell line (clone D42) was generated from an Acly^f/f^ C57BI/6J mouse (Zhao et al., 2016) that had liver tumors induced by a single-administration of diethylnitrosamine (25 mg/kg) via intraperitoneal injection at 2 weeks of age and high-fructose diet (Tekland TD.89247) starting at 6 weeks of age. Following euthanasia at 9 months of age, a palpable tumor was excised from the liver avoiding surrounding tissue. The tumor was digested using Miltenyi Liver Dissociation Kit (mouse 130-105-807), and subject to manual dissociation by pipette to form a single cell suspension in DMEM F-12, 10% heat-inactivated FBS, penicillin/streptomycin, and ITS+ Premix Universal Culture Supplement mix (Corning # 354352). The single cell suspension was seeded onto a collagen-coated 6 cm tissue culture dish. Fibroblasts were depleted by differential trypsinization – cells were trypsinized briefly, and detached cells were removed by gentle rinsing with PBS and aspiration. The remaining adherent cells were trypsinized and reseeded using a limiting dilution to derive a proliferating clonal population. After clonal expansion, cells were maintained in DMEM/F-12 with standard 10% calf serum and then DMEM 10% calf serum. *Acly^-/-^* liver cancer (clone D42C4) cells were generated from Acly^f/f^ liver cancer cells (clone D42) infected with adenoviral Cre recombinase obtained from the University of Pennsylvania Vector Core. Single cell clonal D42 *Acly^-/-^* cell lines were then generated by limiting dilution and ACLY loss was validated by Western blot. Cell lines were generated while cultured in DMEM/F-12 media (Gibco #11320033) supplemented with 10% fetal calf serum and penicillin/streptomycin.

### Human heart tissue

Heart tissue was derived from a male organ donor with no history of heart failure, diabetes mellitus or obesity. All study procedures were approved by the University of Pennsylvania Hospital Institutional Review Board, and prospective informed consent for research use of heart tissue was obtained from organ donor next-of-kin. The heart received *insitu* cold cardioplegia and was placed on wet ice in 4 °C Krebs-Henseleit Buffer. Transmural left ventricular samples, excluding epicardial fat, were cut into 20 mg pieces for and each piece was immediately transferred to a pre-chilled 1 ml Potter-Elvehjem Tissue Grinder (Corning cat. #7725T-1) containing SILEC Hepa1c17 cells in buffer and subjected to fractionation as described below. SILEC Hepa1c17 cells (2.7 E7 cells/sample) were used.

### Liver tissue

All animal studies were carried out in accordance with the IACUC guidelines of the University of Pennsylvania. Male C57Bl6 mice fed ad libitum on a chow diet (Laboratory Autoclavable Rodent Diet 5010, LabDiet cat #0001326) were sacrificed at 5 months old by cervical dislocation at 7 am. The frontal lobe was removed to a dish on ice and a 15 mg piece cut and weighed for fractionation. The weighed piece was immediately transferred to a prechilled 1 ml Potter-Elvehjem Tissue Grinder (Corning cat. #7725T-1) containing SILEC HepG2 cells in fractionation buffer prepared as described below. The equivalent of 110cm dish of 80% confluent SILEC HepG2 cells (~1E7 cells) was used for each sample. Each n represents a different mouse.

### SILEC cell preparation

SILEC labeling of cell lines at an efficiency of >99% across all measurable acyl-CoA species was achieved through the passaging of cells in ^15^N_1_^13^C_3_-pantothenate (Vitamin B5) for at least 9 passages as previously described (Basu et al., 2011). SILEC media was prepared by the addition ^15^N_1_^13^C_3_-pantothenate (Isosciences) (1 mg/L) to custom pantothenate free DMEM (Gibco, Thermo Fisher Scientific) containing 25 mM glucose and 4 mM glutamine with all other components at concentrations according to the standard formulation (Dulbecco and Freeman, 1959). Charcoal:dextran stripped fetal bovine serum (Gemini Biosciences cat. #100-199) was added to 10% (v/v). Labeling efficiency was tested by isotopologue enrichment analysis with comparison to unlabeled control cells. Each batch of charcoal:dextran stripped fetal bovine serum was tested for sufficient labeling efficiency after 3 passages since this can vary from batch to batch(Snyder et al., 2014).

SILEC labeling was performed in multiple cell lines to match experimental cells. To reduce potential for interference from the small fraction of unlabeled acyl-CoA in SILEC cells, and the expense of each experiment, SILEC internal standard was added at lower abundance than experimental cells. However, SILEC internal standard abundance was maintained within the same order of magnitude for analytical robustness. Thus, one third to half the amount of SILEC cells were used for each experimental cell sample.

### Mitochondria and cytosol isolation by differential centrifugation

Mitochondrial and cytosolic fractions were isolated by classical differential centrifugation protocol (Clayton and Shadel, 2014; Frezza et al., 2007), with adaptations for speed and purity (illustrated in **Fig 1C).** SILEC cells matched to the experimental cell type were harvested first. Media was poured from SILEC cell dishes into a waste container and cells were placed on ice at a 45° angle and residual media drained and aspirated completely. Dishes were laid flat on ice and 1 ml ice-cold buffer (210 mM mannitol, 70 mM sucrose, 5 mM Tris-HCl (pH 7.5), 1 mM EDTA (pH 8), adjusted to pH 7.5) added to each dish. Cells were scraped into the buffer, mixed against the plate with a P1000 pipette 6 times to break up cell clumps and combined in a 50 ml tube on ice. The volume was made up to >n*1 ml with buffer (1 ml required per sample). A homogenous cell suspension was made by mixing against the wall of the tube 6 times using a 10 ml pipette immediately before addition to experimental dishes.

For brown adipocytes each sample was a confluent 10 cm dish (~4E5 cells/sample), for *Ady-/-* mouse embryonic fibroblasts each sample was a 80% confluent 15 cm dish (~0.6E5 cells/sample), for HepG2 cells each sample was 1 10 cm dish at 80% confluence (~1E7 cells/sample). Media was removed from experimental cells in the same manner as for the SILEC cells and 1 ml of ice-cold homogenous SILEC cell suspension was added to each. Cells were scraped into the SILEC suspension and transferred to a pre-chilled 1 ml Potter-Elvehjem Tissue Grinder (Corning cat. #7725T-1) in a beaker of ice and water. For standard curve samples, SILEC cell suspension without experimental cells was transferred directly to tissue grinder. Cells were lysed by stroking with the pestle attached to an overhead stirrer (SOS20, Southwest Science) operated at 1,600 rpm. Optimal stroke number was determined for each cell line (10 strokes for Hepa1c17 cells, 15 strokes for HepG2 and liver cancer cell lines (D42/D42C4), 20 strokes for heart tissue and brown adipocytes, 30 strokes for MEF cells) by assessing the purity and integrity of mitochondria and cytosol by Western blot and the intensity of acyl-CoA signal within each compartment across a range up to 60 strokes. Homogenate was transferred to 1.5 ml tubes on ice. For WCL analysis, a 100 μl aliquot of homogenate (representing 10% of the total sample) was removed and quenched in 1 ml ice-cold 10% TCA (Sigma cat. #T6399) in water. Homogenate was centrifuged at 1,300 ×*g* from 10 min at 4 °C and supernatant was transferred to a new pre-chilled 1.5 ml tube. The high-density debris pellet was quenched by resuspension in 1 ml 10% TCA and the supernatant was centrifuged at 10,000 ×g for 20 min at 4 °C to pellet mitochondria. The supernatant (the cytosolic fraction) was quenched by transferal to a new 1.5 ml tube containing 0.25 ml of 50% (w/v) TCA in water to make a final concentration of 10% TCA. Residual cytosolic fraction was carefully removed from the mitochondrial pellet with P200 pipette, and the pellet was quenched by resuspension in 1 ml 10% (w/v) TCA in water. Samples were stored at −80 °C before thawing on ice for acyl-CoA processing, or directly processed.

### Nuclear isolation

Nuclear isolation was achieved by detergent assisted hypoosmotic lysis and differential centrifugation (illustrated in **Figure 5A).** SILEC cells matched to the experimental cell type were harvested before experimental cells. Media was poured from SILEC cell dishes into a waste container, and cells were placed on ice at a 45° angle and residual media drained and aspirated completely. Dishes were laid flat on ice and 0.5 ml ice-cold lysis buffer (250 mM sucrose, 15 mM Tris-HCl (pH 7.5), 60 mM KCl, 15 mM NaCl, 5 mM MgCI_2_, 1 mM CaCl_2_ adjusted to pH 7.4) added to each dish. SILEC cells were scraped into the buffer, cell clumps were broken up by pipetting up and down against the plate 4 times with a P1000. Cells were then combined in a 15 ml tube on ice. The volume was made up to > 0.5*n ml (0.5 ml required per sample) with buffer and kept on ice. Immediately before addition to experimental dishes, NP-40 (1% v/v in lysis buffer) was added to achieve a final concentration of 0.1% (v/v) and the cell suspension was homogenized by mixing by laminar flow against the wall of the 15 ml tube 4 times with a 5 ml pipette with care taken to avoid frothing. NP-40 addition was delayed to coordinate detergent lysis of SILEC cells with experimental cells.

Media was removed from experimental cells in the same manner as for the SILEC cells and 0.5 ml of ice-cold homogenous SILEC cell suspension was added to each dish/sample. Cells were scraped into the SILEC suspension, mixed against the plate with a P1000 pipette 4 times to break up cell clumps and transferred to a 1.5 ml tube on ice. For WCL analysis, 50 μl (representing 10% of the total sample) was removed after homogenization and quenched in 1 ml ice-cold 10% TCA in water. Nuclei were pelleted by centrifugation at 600 ×g for 5 min at 4 °C. The supernatant (the ‘non-nuclear’ fraction) was quenched by transferal to a new 1.5 ml tube containing 0.125 ml of 50% (w/v) TCA in water to make a final concentration of 10% TCA. The nuclear pellet was washed by the addition of 0.5 ml lysis buffer without NP-40 and recentrifuged at 600 *×g* for 5 min at 4 °C. The supernatant (the ‘wash’ fraction) was quenched by transferal to a new 1.5 ml tube containing 0.125 ml of 50% (w/v) TCA in water to make a final concentration of 10% TCA. Residual wash was carefully removed from the nuclear pellet with a P200 pipette, and the nuclear pellet was quenched by the resuspension in 1 ml 10% w/v TCA in water.

### Standard curve generation

Separate standard curves were generated for each sub-cellular fraction. Equal aliquots of SILEC internal standard cells were fractionated in the absence of experimental unlabeled cells and known quantities of unlabeled standards were added before extraction **(Supp 1A).** 6-point standard curves (standard 0 to standard 5) were generated for each fraction by dilution from a stock mixture of unlabeled acyl-CoA standards. The stock mixture was dissolved in 10% TCA in water according to **Table 2** and aliquots were frozen at −80 and thawed once at use. Serial 3-fold dilutions were made from standard 5 stock in 10% TCA in water and each dilution was added in a volume of 100 μl to the appropriate samples containing SILEC internal standard. The concentration of standard 5 (the highest standard) stock was adjusted to suit metabolite abundance in different samples. For most cell samples standard 5 stock was a 50x dilution of the original stock mixture (mitochondrial and nuclear fractions used a 250x dilution). For most tissue samples, standard 5 stock was a 20x dilution of stock mixture (mitochondrial and nuclear fractions used a 100x dilution).

Standard curves were generated by plotting standard quantity against relative intensity (light/ SILEC internal standard) and linear curves were fitted (formula 1). This standard curve was used to determine the quantity recovered within each sample (formula 2). Adjustments were made to account for removal of 10% of sample for whole cell lysate aliquot and for cell number before fractionation with formula 3 for whole cell lysate and formula 4 for all other fractions.

Where *y* = relative intensity, *x_s_* = standard quantity, *b* = constant, *x_Q_* = quantity recovered in sample, *C* = cell number before fractionation, Q = adjusted quantity in sample.

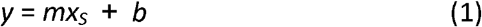

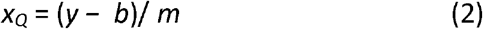

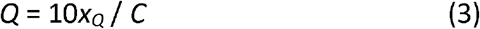

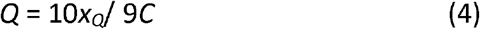

### SILEC-SF Mito-IP

Mito-IP was performed as described (Chen et al., 2017) with adjustments to accommodate SILEC internal standard addition and acyl-CoA extraction in TCA. Breifly, HepG2 cells were infected with retrovirus generated in Pheonix-AMPHO cells transfected with pMXS-3XHA-EGFP-OMP25 construct. Following blasticidin selection (5 ug/ml) Mito tag expressing cells were sorted into 3 categories (low/no expression, medium and high) based on GFP expression level using a BD FACS Jazz cell sorter. Medium expression cells were cultured for further use. SILEC cells were generated by passaging these cells in SILEC media as described above. Cells were harvested by addition of SILEC cells, and homogenized as described above for mitochondria and cytosol isolation by differential centrifugation except that the buffer used was KPBS (136 mM KCl and 10 mM KH2PO4, pH 7.25). Each sample was 1 15 cm dish at 80% confluence (~1E7 cells/sample). Homogenate was transferred to a 1.5 ml tube on ice and centrifuged at 1,000 xg for 2 min at 4 °C. The post-nuclear supernatant was transferred to a 1.5 ml tube containing 200 ul of prewashed Anti-HA magnetic beads and incubated by rotation for 3.5 min at 4 °C. Beads were collected on a magnet and the overlying solution ‘mitochondria depleted cytoplasm’ was transferred to a 1.5 ml and quenched by addition of TCA to a final concentration of 10% (w/v). Beads were washed twice with 1 ml KPBS before extraction of mitochondria by addition of TCA 10% w/v. Samples were incubate on ice for 10 min and vortexed vigorously to extract metabolites. The supernatant was then removed from the beads for further processing.

### SILEC-SF Mito-IP standard curve generation

For Mito-IP, standard curves were generated separately for each experimental group in each fraction by standard addition **(Figure S1B).** Following sonication, half of each replicate sample was pooled together, vortexed mixed and then redivided into 3 exact replicates. 3 dilutions of unlabeled acyl-CoA standard stock mixture **(Table 2)** were made in 10% TCA. For non-mitochondrial fractions the dilutions were (50x, 150x, 450x), for mitochondria (500x, 1500x, 4500x). Each dilution was added in a volume of 100 μl to a replicate sample and samples were processed and analyzed by LC-MS. Standard curves were generated by plotting the known concentration of acyl-CoA standard stock against the relative signal intensity (unlabeled/SILEC internal standard) and fitting a curve by linear regression **(Figure S1B).** The linear equation (y = mx + b, where y = relative intensity and x = quantity) was adjusted to account for the presence of additional unlabeled analyte in the standard curve derived from the sample (y = mx). This formula was then used to calculate quantity within each of the replicate samples to which no standard was added. Values were doubled to account for half the sample being used in the standard curve.

### Normalization

Data were normalized to cell counts from an extra replicate cell dish for each condition. Cells were trypsinzed and total cell number and volume was determined by Coulter counter (Coulter), cell number was also determined manually by counting using a haemocytometer. For tissue samples, tissue pieces were weighed on a balance to +/- 0.01 mg before fractionation and data was normalized to mass for each sample.

### Whole cell direct extraction

Media was poured from dishes into a waste container and cells were placed on ice at a 45° angle and residual media aspirated completely. Dishes were laid flat on ice and 1 ml 10% TCA in water was added. Internal standard extracted from yeast grown in ^15^N_1_^13^C_3_-pantothenate as previously described (Snyder et al., 2015) was added (100 μl/sample). Cells were scraped into the TCA and transferred to 1.5 ml tubes on ice then either processed directly or stored at −80 °C. Standard curves were generated in parallel with equal aliquots of internal standard in 10% TCA in water.

### Stable isotope tracing in whole cells

For experiments involving 50% diluted isoleucine, valine, or leucine tracer, cells were preincubated in serum-free DMEM (Thermo Fisher Scientific, Gibco #11965084) for 8 hours before media was exchanged for tracing media. Cells were incubated in tracing media at 37 °C and 5% CO_2_ for 18 hours before whole cell direct extraction. Tracing media was prepared using DMEM base lacking glucose or glutamine (Thermo-Fisher scientific, Gibco cat. #A1443001). Glucose was replaced at (4.5 g/L), and glutamine at (584 mg/L) and additional isoleucine (105 mg/L), valine (94 mg/L) and leucine (105 mg/L) were added. Except for the targeted experimental tracer (U^13^C, see **Table** 1), all additives were unlabeled. Total substrate concentrations were equal across all samples except propionate, which was added only to the U^13^C-propionate tracing samples at 1 mM.

**TABLE 1:**
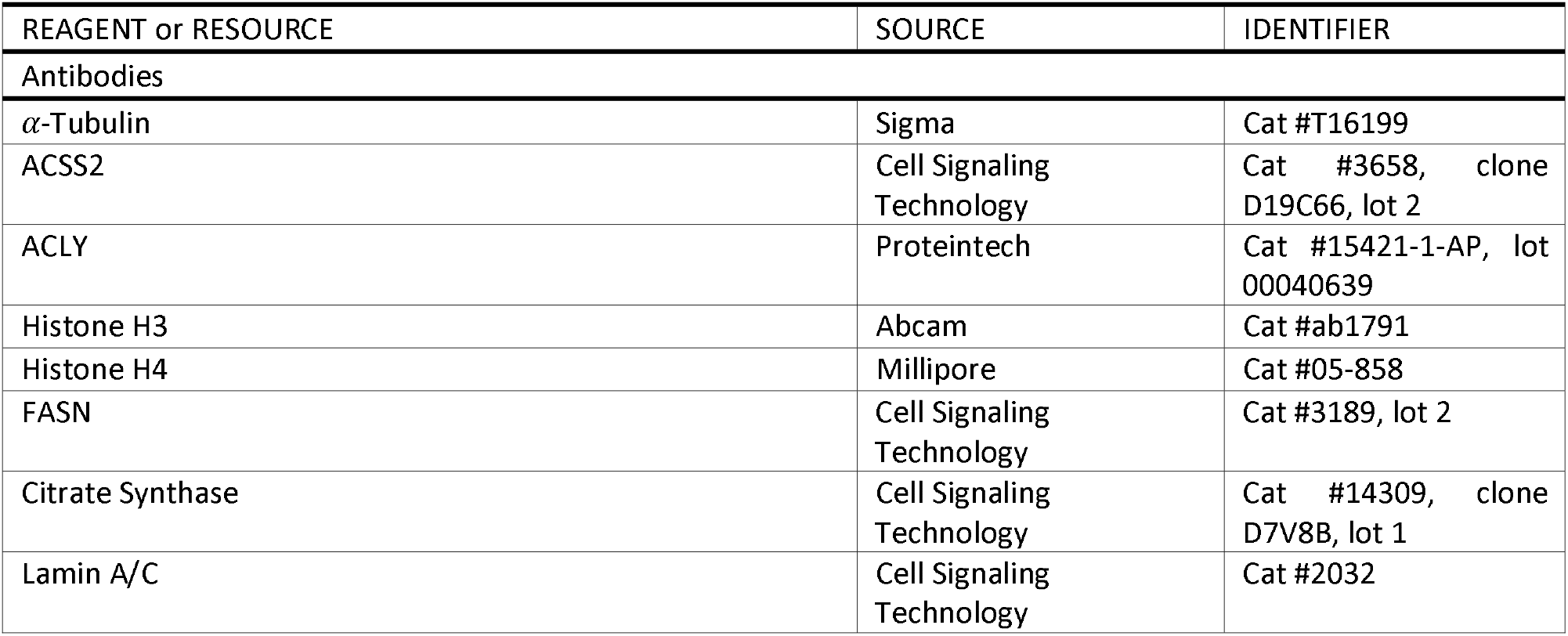
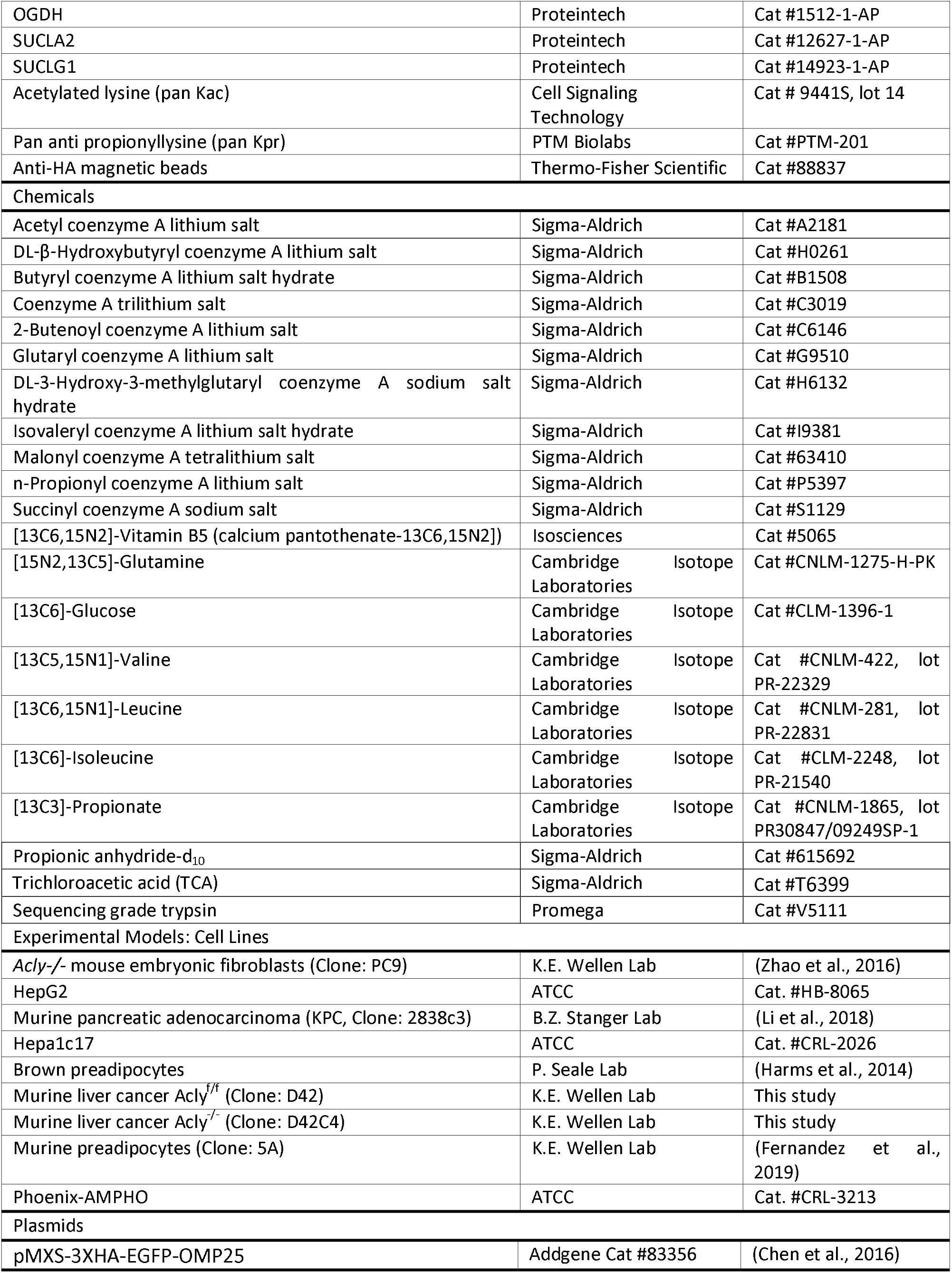
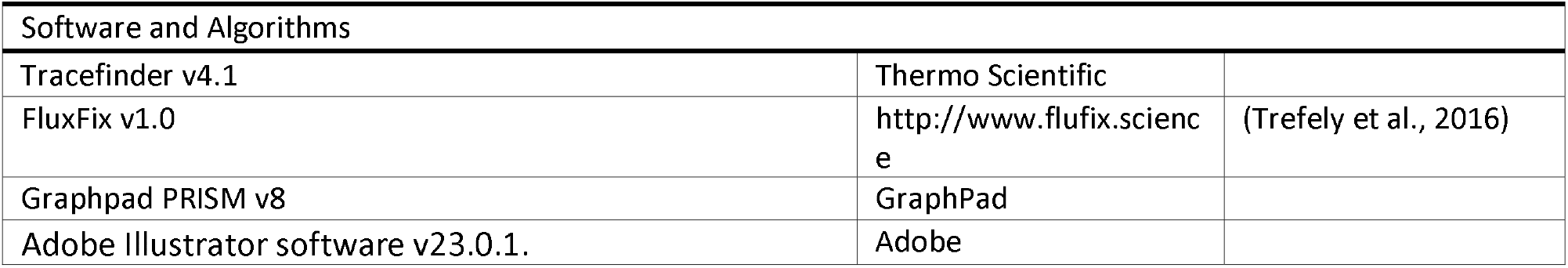
KEY RESOURCES.

**Table 2:** Acyl-CoA standard mixture.

| Unlabeled standard | Concentration (pmol/100 μl) |
| --- | --- |
| Acetyl-CoA | 5000 |
| BHB-CoA | 500 |
| Butyryl-CoA | 500 |
| CoASH | 500 |
| Crotonoyl-CoA (2-butenoyl-CoA) | 500 |
| Glutaryl-CoA | 500 |
| HMG-CoA | 500 |
| Isovaleryl-CoA | 500 |
| Malonyl-CoA | 500 |
| Propionyl-CoA | 500 |
| Succinyl-CoA | 5000 |

For experiments with undiluted isoleucine labeling, cells were washed with PBS and tracing media was added before incubation at 37 °C and 5% CO_2_. Tracing media was prepared from DMEM base lacking Amino Acids, Glucose, Pyruvic Acid and Phenol Red (Biological Life Science D9800-26) reconstituted with glucose (4.5 g/L), glutamine (2 mM), and other unlabeled amino acids according to DMEM standard formulation(Dulbecco and Freeman, 1959) except for isoleucine which was added as ^13^C_6_-isoleucine (**Table** 1) to (105 mg/L). NaOH and HEPES (25 mM) was added to adjust to pH7.4 and dialyzed fetal bovine serum (Gemini Biosceinces Cat. # 100-108) was added to 10% (v/v).

### Isoleucine and valine withdrawal

Cells were washed with PBS and media replaced with media prepared from DMEM base lacking amino acids, glucose, pyruvic acid and phenol red (Biological Life Science D9800-26) reconstituted with glucose (4.5 g/L), glutamine (2 mM), and other amino acids except for isoleucine and valine according to DMEM standard formulation(Dulbecco and Freeman, 1959). NaOH and HEPES (25 mM) was added to adjust to pH7.4 and dialyzed fetal bovine serum (Gemini Biosceinces Cat. # 100-108) was added to 10% (v/v).

### Sub-cellular stable isotope tracing with post-labeling

Murine pancreatic ductal adenocarcinoma (KPC) cells were incubated with U^13^C-isoleucine tracer before fractionation as described above. Fractionation with post-labeling was carried out as previously described (Trefely et al., 2019) with the addition of U^13^C-isoleucine (525 mg/L) and unlabeled isoleucine (525 mg/L) to fractionation buffer.

### Acyl-CoA sample processing

Samples were thawed and kept on ice throughout processing. Cell and fraction samples in 10% (w/v) trichloroacetic acid (Sigma cat. #T6399) in water were sonicated for 12 × 0.5 s pulses, protein was pelleted by centrifugation at 17,000 *×g* from 10 min at 4 °C. The supernatant was purified by solid-phase extraction using Oasis HLB 1cc (30 mg) SPE columns (Waters). Columns were washed with 1 mL methanol, equilibrated with 1 mL water, loaded with supernatant, desalted with 1 mL water, and eluted with 1 mL methanol containing 25 mM ammonium acetate. The purified extracts were evaporated to dryness under nitrogen then resuspended in 55 μl 5% (w/v) 5-sulfosalicylic acid in water.

### Acyl-CoA analysis by LC-MS

Acyl-CoAs were measured by liquid chromatography-high resolution mass spectrometry. Briefly, 5-10 μl of purified samples in 5% SSA were analyzed by injection of an Ultimate 3000 Quaternary UHPLC coupled to a Q Exactive Plus (Thermo Scientific) mass spectrometer in positive ESI mode using the settings described previously (Frey et al., 2016). Quantification of acyl-CoAs was via their MS2 fragments and the targeted masses used for isotopologue analysis are indicated in **Table 3.** Data were integrated using Tracefinder v4.1 (Thermo Scientific) software. Isotopic enrichment in tracing experiments was calculated by normalization to unlabeled control samples using the FluxFix calculator (Trefely et al., 2016).

**Table 3:**
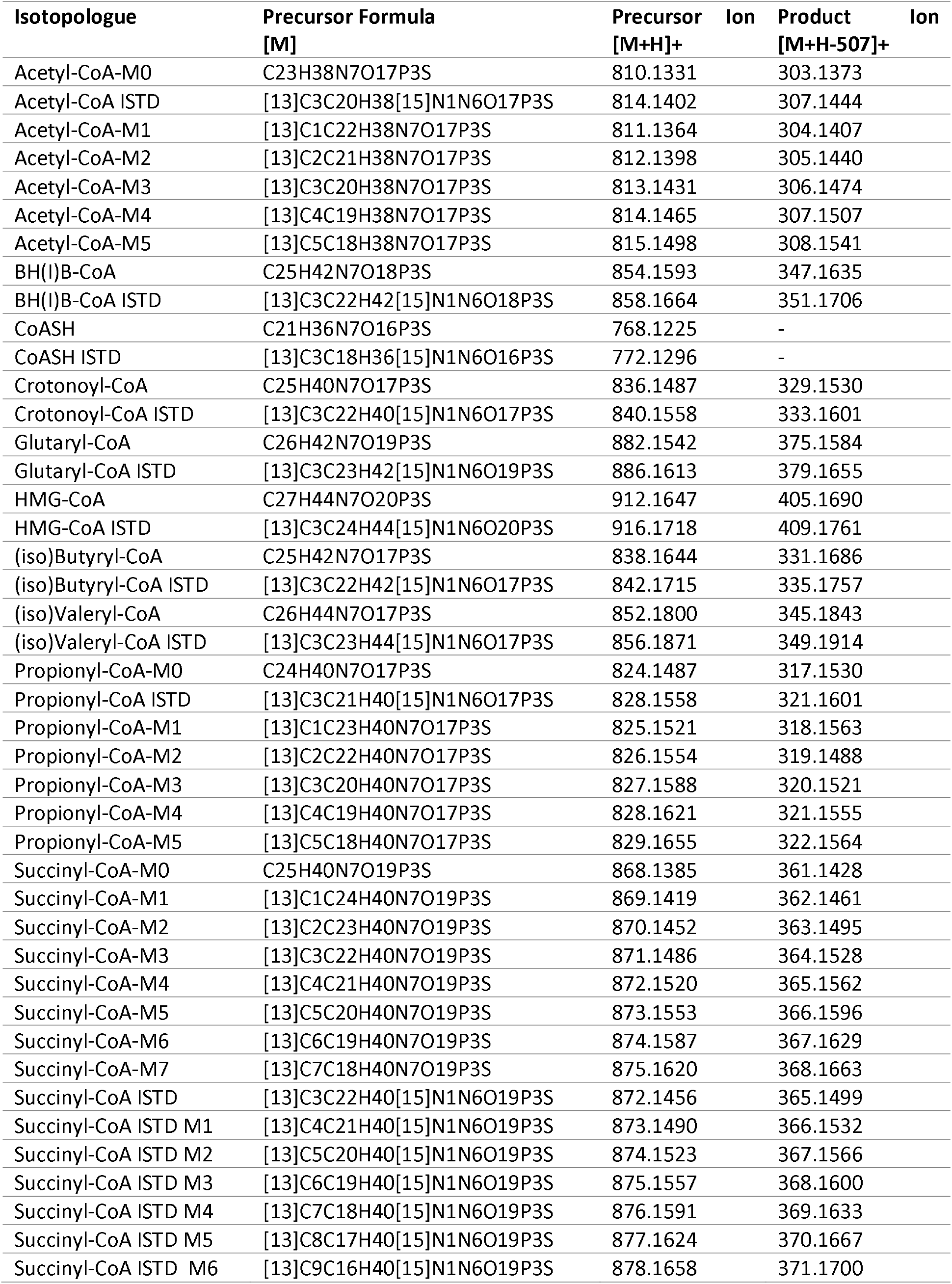
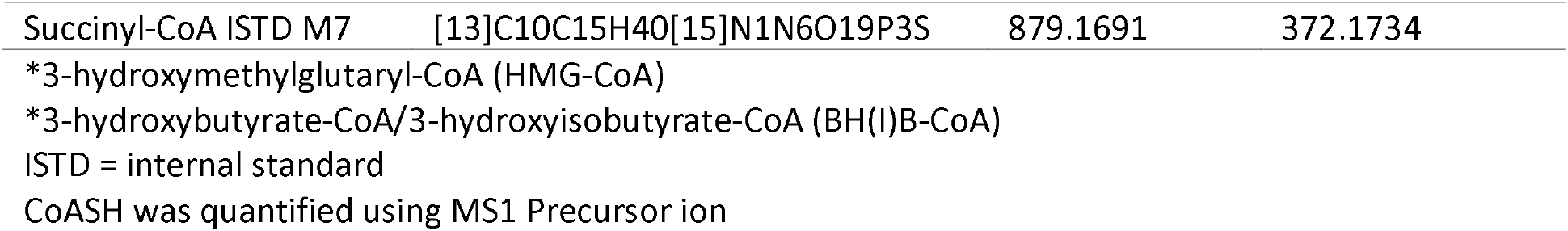
Acyl-CoA Masses.

### Western blotting

Western blotting was performed using mini gel tank system (Life Biotechnologies) with 4-12% gradient Bis-Tris gels (NuPage, Invitrogen cat. #NP0335) and 0.45 μm pore size nitrocellulose membranes (BioRad cat. #1620115) or 0.22 μm pore size for histone blots. Membranes were probed with antibody (see **Table** 1) according to the manufacturer’s instructions. An Odyssey CLx imaging system with Image Studio v2.0.38 software (LI-COR Biosciences) was used to acquire images which were exported as TIFF files then cropped and arranged using Adobe Illustrator software v23.0.1.

### Histone acid extraction

Histones were acid-extracted after cell treatment as previously described (Sidoli et al., 2016). For each sample, a 10 cm dish of 80% confluent cells was washed with ice cold PBS on ice. Cells were scraped in 0.25 ml of NIB-250 buffer (15 mM Tris-HCl (pH 7.5), 60 mM KCl, 15 mM NaCl, 5 mM MgCl_2_, 1 mM CaCl_2_, 250 mM sucrose, 1 mM DTT, 10 mM sodium butyrate, 1x protease inhibitor cocktail (Sigma-Aldrich P8340)) + 0.1% NP-40. Nuclei were pelleted by centrifugation at 600 xg for 5 min at 4°C. The pellet was then washed twice by addition of 0.25 ml NIB-250 buffer without NP40, centrifugation at 600 xg, and removal of supernatant. Nuclear pellet was resuspended in 0.6 ml of 0.4N H_2_SO_4_ and extraction with rotation at 4°C for 16 h. Centrifugation at 11,000 xg for 10 min at 4°C sedimented precipitate. The supernatant was taken to a new 1.5 ml tube and histones were precipitated on ice for 16 h by addition of 100% (w/v) trichloroacetic acid to a final concentration of 20%. The histone pellet was sedimented by centrifugation at 11,000 xg for 10 min at 4°C. Supernatant was removed and the histone pellet was washed with 1 ml acetone with 0.1% 12N HCl and centrifuged at 11,000 xg for 10 min at 4°C. Pellet was washed again with 1 ml acetone and air dried.

### Histone acyl proteomic analysis

#### Histone digestion

Acid extracted histone pellets were dissolved in 50 mM ammonium bicarbonate, pH 8.0, and histones were subjected to derivatization with as previously described (Sidoli et al., 2019b). Briefly, intact histones were mixed with 5 μL of deuterium-labeled propionic anhydride (propionic anhydride-d_10_) rapidly followed by 14 μL of ammonium hydroxide (Sigma-Aldrich) to adjust to pH 8.0. The mixture was incubated for 15 min and the procedure was repeated. Histones were then digested with 1 μg of sequencing grade trypsin (Promega) diluted in 50 mM ammonium bicarbonate (1:20, enzyme:sample) overnight at room temperature. Derivatization reaction was repeated to derivatize peptide N-termini. The samples were dried in a vacuum centrifuge.

Prior to mass spectrometry analysis, samples were desalted using a 96-well plate filter (Orochem) packed with 1 mg of Oasis HLB C-18 resin (Waters). Briefly, the samples were resuspended in 100 μl of 0.1% TFA and loaded onto the HLB resin, which was previously equilibrated using 100 μl of the same buffer. After washing with 100 μl of 0.1% TFA, the samples were eluted with a buffer containing 70 μl of 60% acetonitrile and 0.1% TFA and then dried in a vacuum centrifuge.

#### LC-MS/MS Acquisition and Analysis

Samples were resuspended in 10 μl of 0.1% TFA and loaded onto a Dionex RSLC Ultimate 3000 (Thermo Scientific), coupled online with an Orbitrap Fusion Lumos (Thermo Scientific). Chromatographic separation was performed with a two-column system, consisting of a C-18 trap cartridge (300 μm ID, 5 mm length) and a picofrit analytical column (75 μm ID, 25 cm length) packed in-house with reversed-phase Repro-Sil Pur C18-AQ 3 μm resin. Histone peptides were separated using a 30 min gradient from 1-30% buffer B (buffer A: 0.1% formic acid, buffer B: 80% acetonitrile + 0.1% formic acid) at a flow rate of 300 nl/min. The mass spectrometer was set to acquire spectra in a data-independent acquisition (DIA) mode using isolation windows as previously described (Sidoli et al., 2015). Briefly, the full MS scan was set to 300-1100 m/z in the orbitrap with a resolution of 120,000 (at 200 m/z) and an AGC target of 5×10e5. MS/MS was performed in the orbitrap with sequential isolation windows of 50 m/z with an AGC target of 2×10e5 and an HCD collision energy of 30.

Histone peptides raw files were imported into EpiProfile 2.0 software (Yuan et al., 2018). From the extracted ion chromatogram, the area under the curve was obtained and used to estimate the abundance of each peptide. In order to achieve the relative abundance of post-translational modifications (PTMs), the sum of all different modified forms of a histone peptide was considered as 100% and the area of the particular peptide was divided by the total area for that histone peptide in all of its modified forms. The relative ratio of two isobaric forms was estimated by averaging the ratio for each fragment ion with different mass between the two species. The resulting peptide lists generated by EpiProfile were exported to Microsoft Excel and further processed for a detailed analysis. To assess the incorporation rate of the ^13^C_3_-Kpr, i.e. histone propionylation, we performed manual signal extraction using Xcalibur QualBrowser (Thermo) and the area under the curve was used as representative of the peptide abundance.

#### Mass spectrometry raw data availability

All raw mass spectrometry data files from histone acyl-proteomic analysis in this study have been submitted to the Chorus repository (https://chorusproject.org/pages/index.html) under project number 1724.

### Graphing and statistical analyses

Data presented are shown either of mean ± standard deviation or, for curve fits, mean ±95% confidence intervals. Graphpad Prism software (v.8) was used for graphing and statistical analysis. For comparison between two groups, datasets were analyzed by two-tailed Student’s ř-test with Welch’s correction and statistical significance defined as p<0.05 (*), p < 0.01 (**), p < 0.001 (***), p < 0.0001 (****).

## Author contributions

ST, NWS, and KEW conceptualized the study and designed experiments. ST prepared figures and wrote the manuscript. NWS and KEW edited the manuscript. ST performed the majority of the experiments and data analysis. JL, KH, MN and CDL performed experiments and analysis. JS, EVK, MD, HJ, AB, HLP and JPZ performed metabolite extraction and analysis. SS and SS performed histone acyl proteomic analyses. LI and SZ generated D42 and D42C4 liver cancer cell lines. JER and KCB procured heart samples. CM provided valuable advice and support with mass spectrometry. JB-S provided useful discussion. All authors read and provided feedback on manuscript and figures.

## Acknowledgements

NWS was supported by R01GM132261 and P30ES013508. KEW is supported by R01CA228339, R01DK116005, R01CA174761 and R01CA248315. ST was supported by the American Diabetes Association through post-doctoral fellowship 1-18-PDF-144. KH and JB-S were supported by the Austrian Science Fund grants FWF W1226and FWF P27108. CDL is supported by NIH T32 GM07170. LI was supported by T32 GM-07229 and 2-T32-CA-115299-13. SZ was supported by F99CA222741. SS and SS are supported by the Leukemia Research Foundation (Hollis Brownstein New Investigator Research Grant), AFAR (Sagol Network GerOmic Award), Deerfield (Xseed award), the NIH P30 grant CA01333047, and the shared instrument grant NIH 1 S10 OD030286-01. We thank Dr. Kenneth B Margulies and the Gift of Life organization for allowing the use of human tissue for this project.

---

**Supplemental Figure 1.**
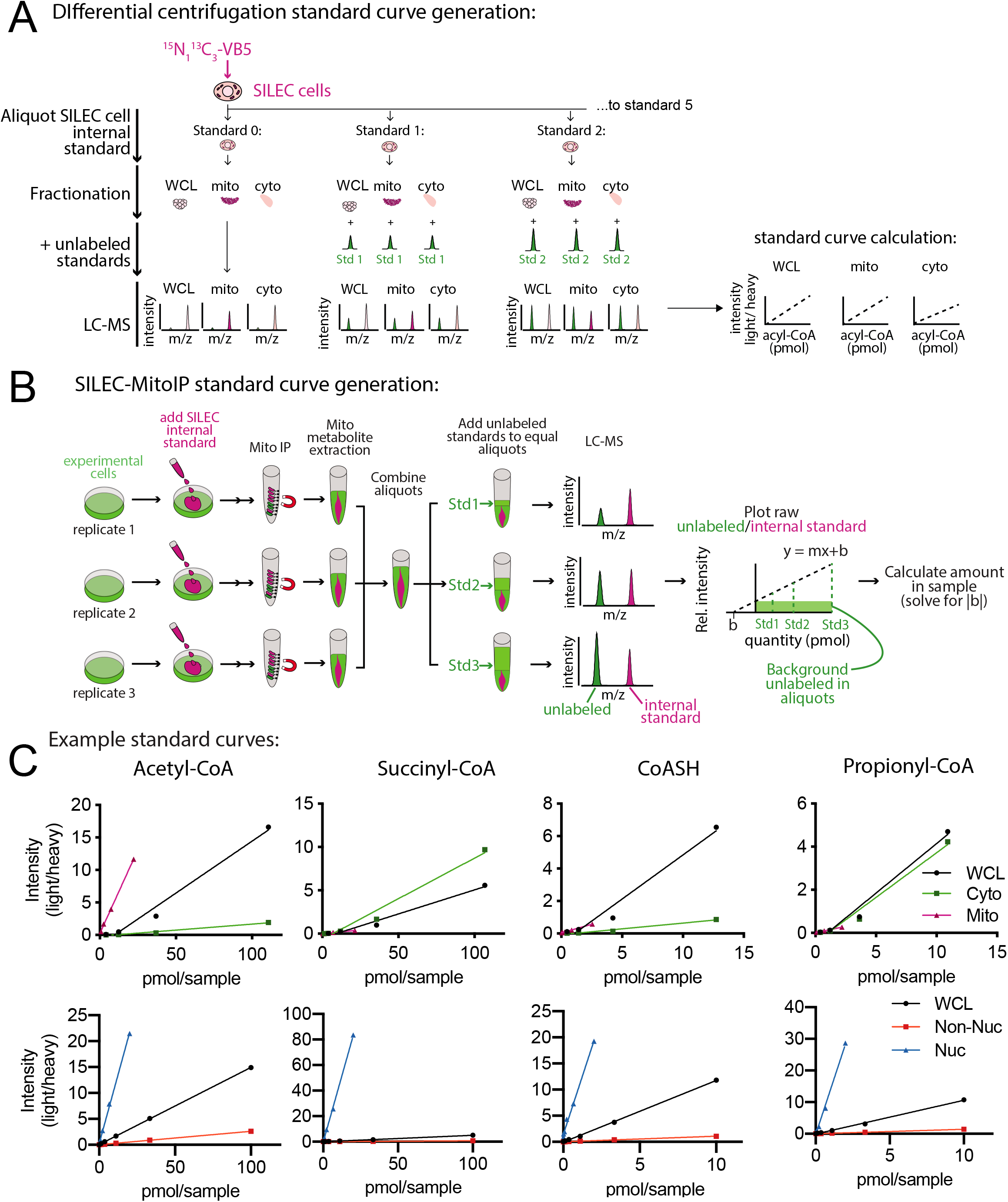
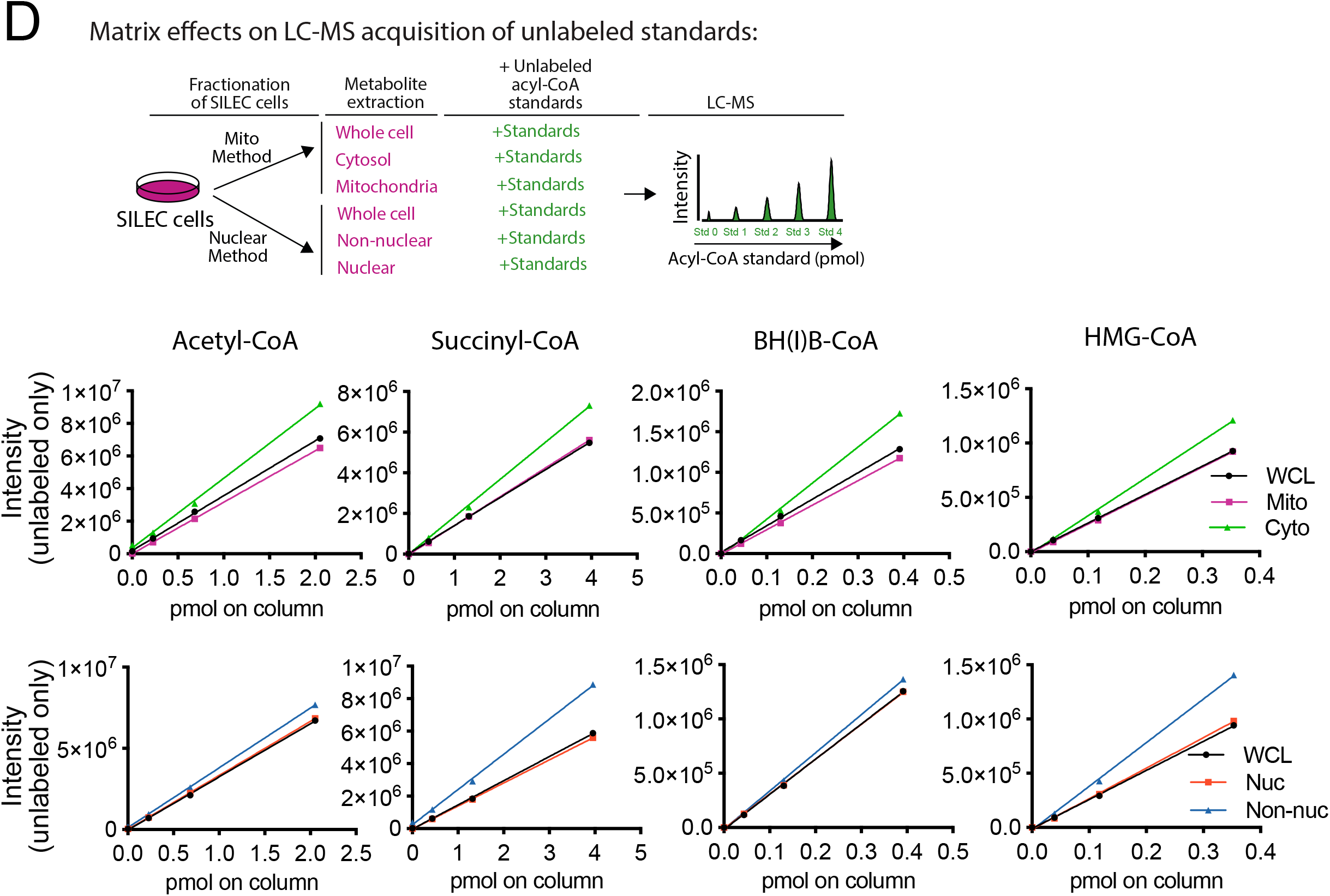
**A)** Schematic representation of standard curve generation in differential centrifugation protocols. Sub-cellular fractions were generated from equal aliquots of SILEC internal standard cells processed in parallel to the experimental + SILEC cell samples. After fractionation, known quantities of unlabeled standards were added to build standard curves. B) Schematic representation of standard curve generation by standard addition in Mito-IP protocol **C)** Representative standard curves generated in different subcellular matrices across several acyl-CoA species in HepG2 cells using differential centrifugation for mitochondrial/cytosol (upper panel) or nuclear (lower panel) isolation. WCL = whole cell lysate. D) Demonstration of subcellular matrix specific effects on acyl-CoA acquisition. Raw signal intensity for unlabeled acyl-CoAs are displayed across a range of concentrations in the presence of different sub-cellular matrices generated from fully labeled Hepa1c17 SILEC cells separated into different fractions.

**Supplemental Figure 2.**
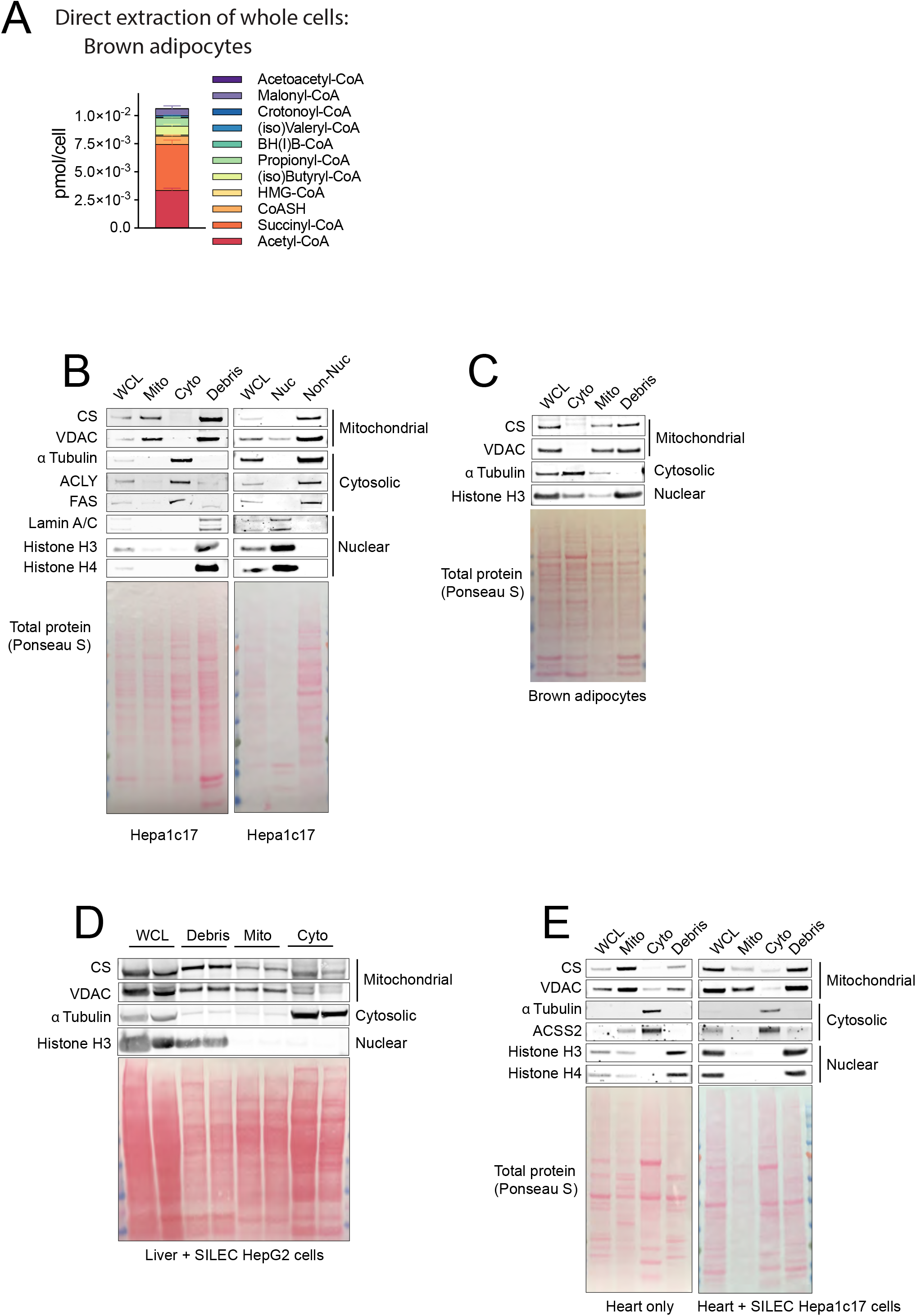
A) Direct extraction of whole brown adipocytes relating to Fig 1C B-E) Western blot analysis of protein distribution. Equal protein quantity was loaded for each fraction. B) Hepa1c17 fractionation C) Brown adipocytes were fractionated using different numbers of strokes for optimization. 20 strokes was used for experiments. D) Liver tissue combined with SILEC HepG2 cells. E) Human heart fractionation for heart only, and heart combined with SILEC Hepa1c17 cell internal standard. Abbreviations: CS (citrate synthase), VDAC (voltage dependent anion channel), FAS (fatty acid synthase), ACSS2 (Acyl-CoA synthetase short chain family member 2), ACLY (ATP-citrate lyase).

**Supplemental Figure 3.**
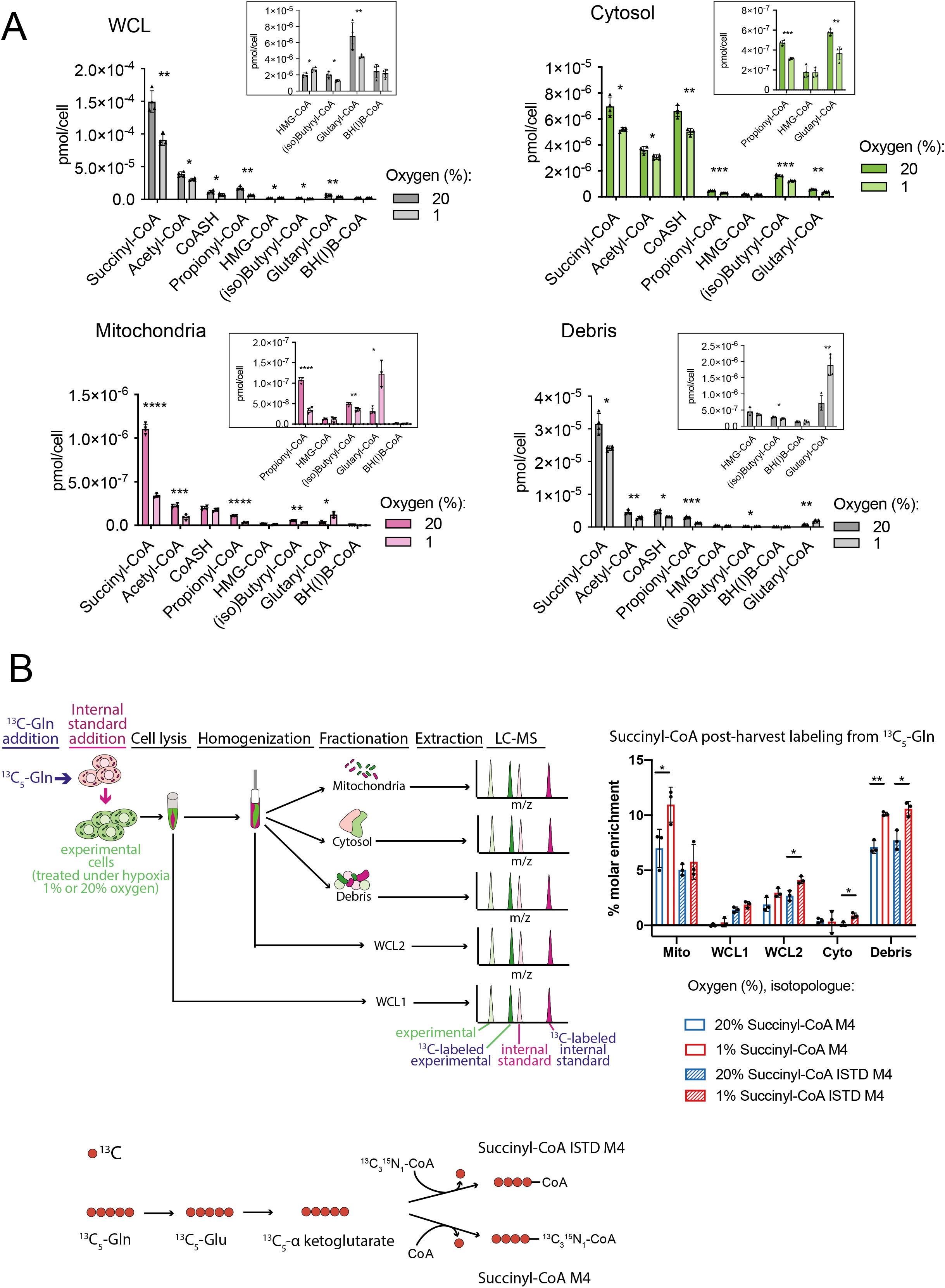
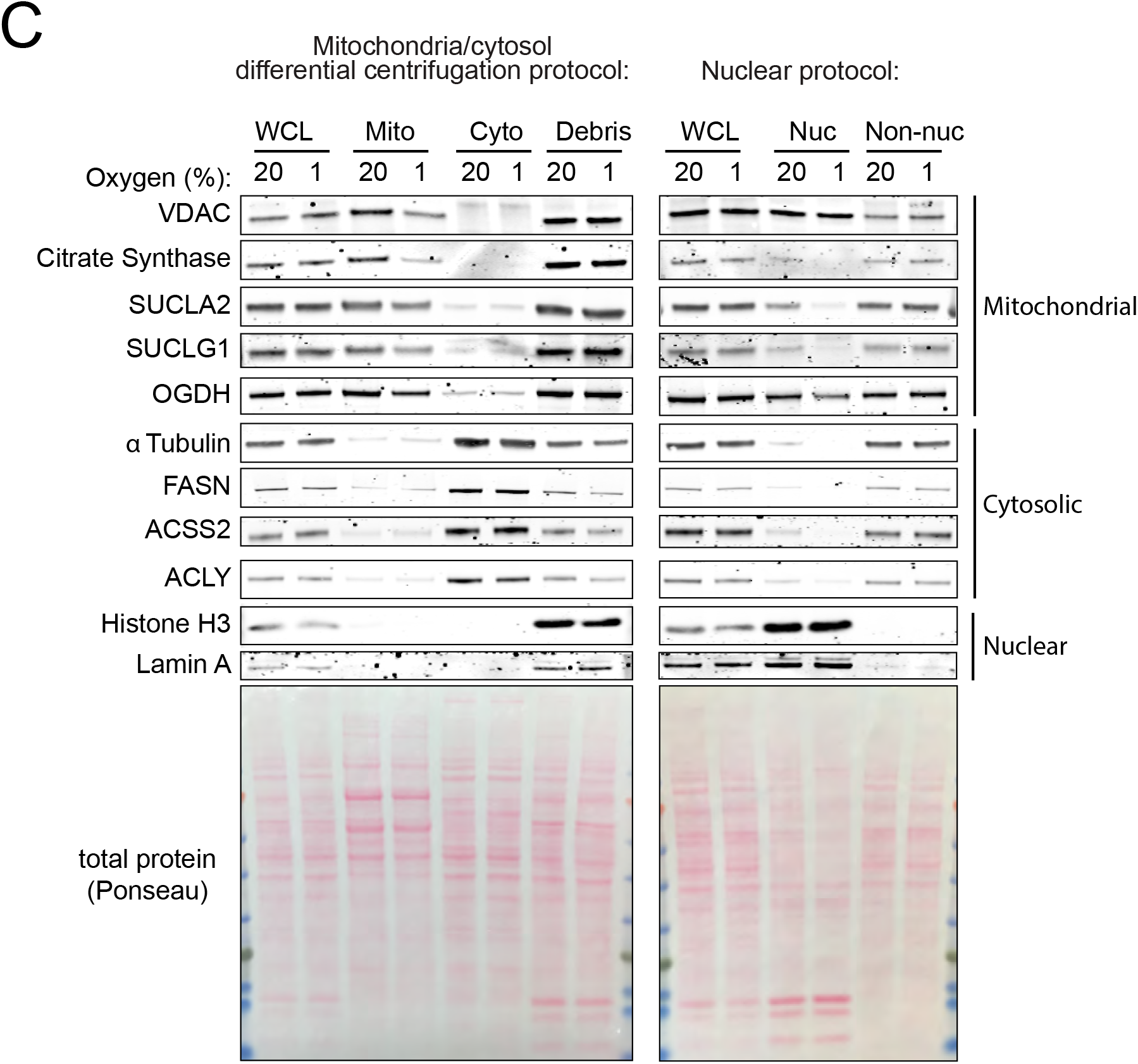
HepG2 cells were incubated under 20% (normoxia) or 1% (hypoxia) for 24 h. Cells were serum starved in DMEM containing 5 mM glucose and 2 mM glutamine under their respective oxygen tensions for 2 h before harvest. **A)** Data relating to **Fig 3C** showing complete set of short chain acyl-CoA species quantified by SILEC-SF using mitochondrial/cytosolic differential centrifugation procedure from a representative experiment, which is also incorporated into **Fig 3D** and **Fig 5E.** Symbols represent individual replicate dishes (n=4) and error bars represent standard deviation. Lower abundance metabolites are magnified in the upper right corner of each panel. B) Post-harvest labeling with ^13^C_5_-Gln was applied to SILEC-SF differential centrifugation method. ^13^C incorporation into the acyl chain of acyl-CoAs derived from experimental cells and SILEC internal standard labeled cells can be differentiated by mass. Postharvest C_5_-Gln incorporation into succinyl-CoA M4 was compared between experimental and SILEC internal standard succinyl-CoA across 2 experimental conditions **C)** Western blots comparing protein enrichment for representative marker proteins for mitochondria, cytosol and nucleus. HepG2 cells were subject to sub-cellular fractionation by both the mitochondrial/cytosol and nuclear/non-nuclear protocols. Equal protein quantity was loaded for each fraction. Abbreviations: VDAC (voltage dependent anion channel), SUCLA2 (Succinyl-CoA ligase [ADP-Forming] subunit beta), SUCLG1 (Succinyl-CoA ligase [GDP-forming] subunit alpha), OGDH (oxoglutarate dehydrogenase), FAS (fatty acid synthase), ACSS2 (Acyl-CoA synthetase short chain family member 2), ACLY (ATP-citrate lyase). For comparison between two groups, datasets were analyzed by two-tailed Student’s ř-test with Welch’s correction and statistical significance defined as p < 0.05 (*), p < 0.01 (**), p < 0.001 (***), p < 0.0001 (****).

**Supplemental Figure 4.**
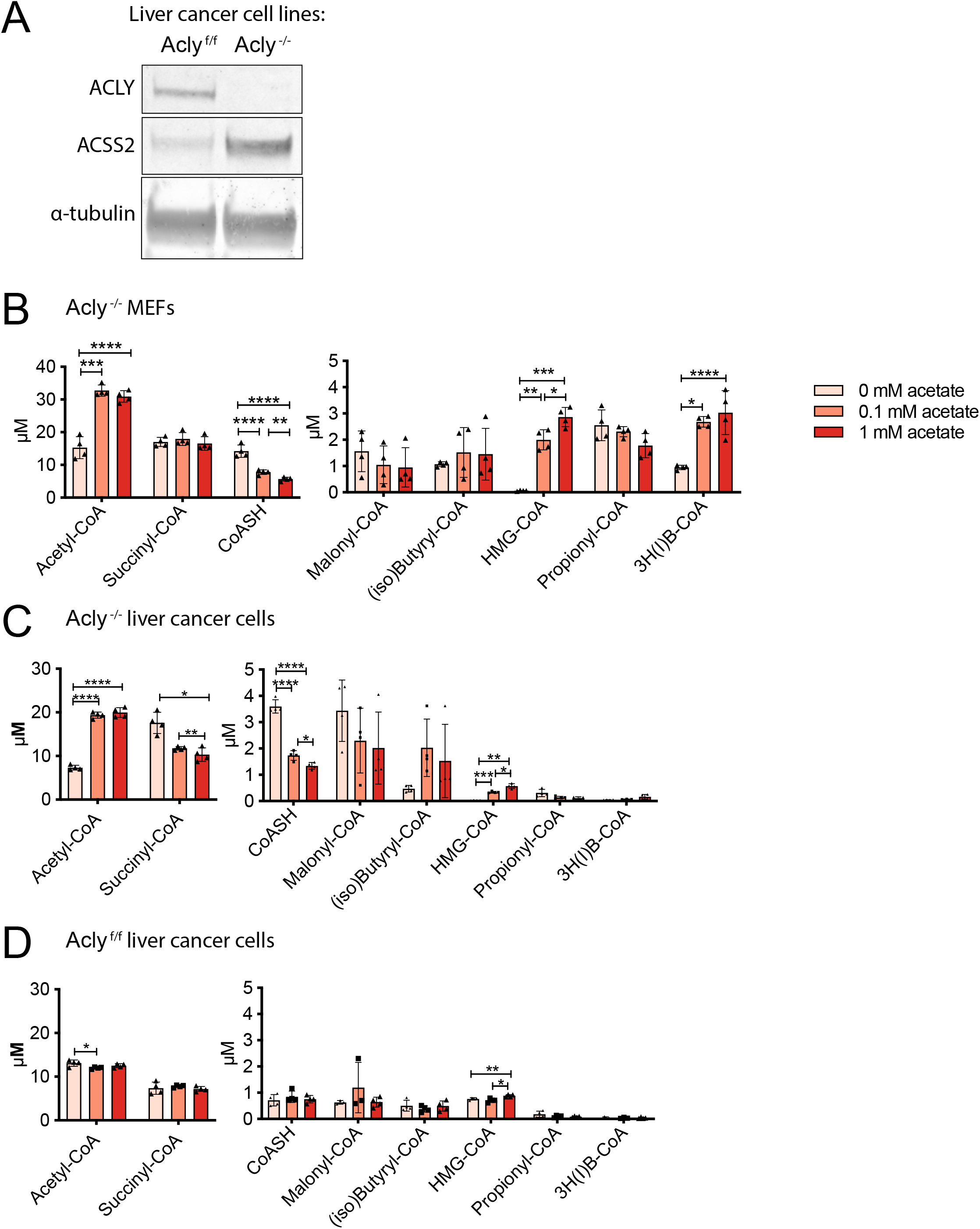
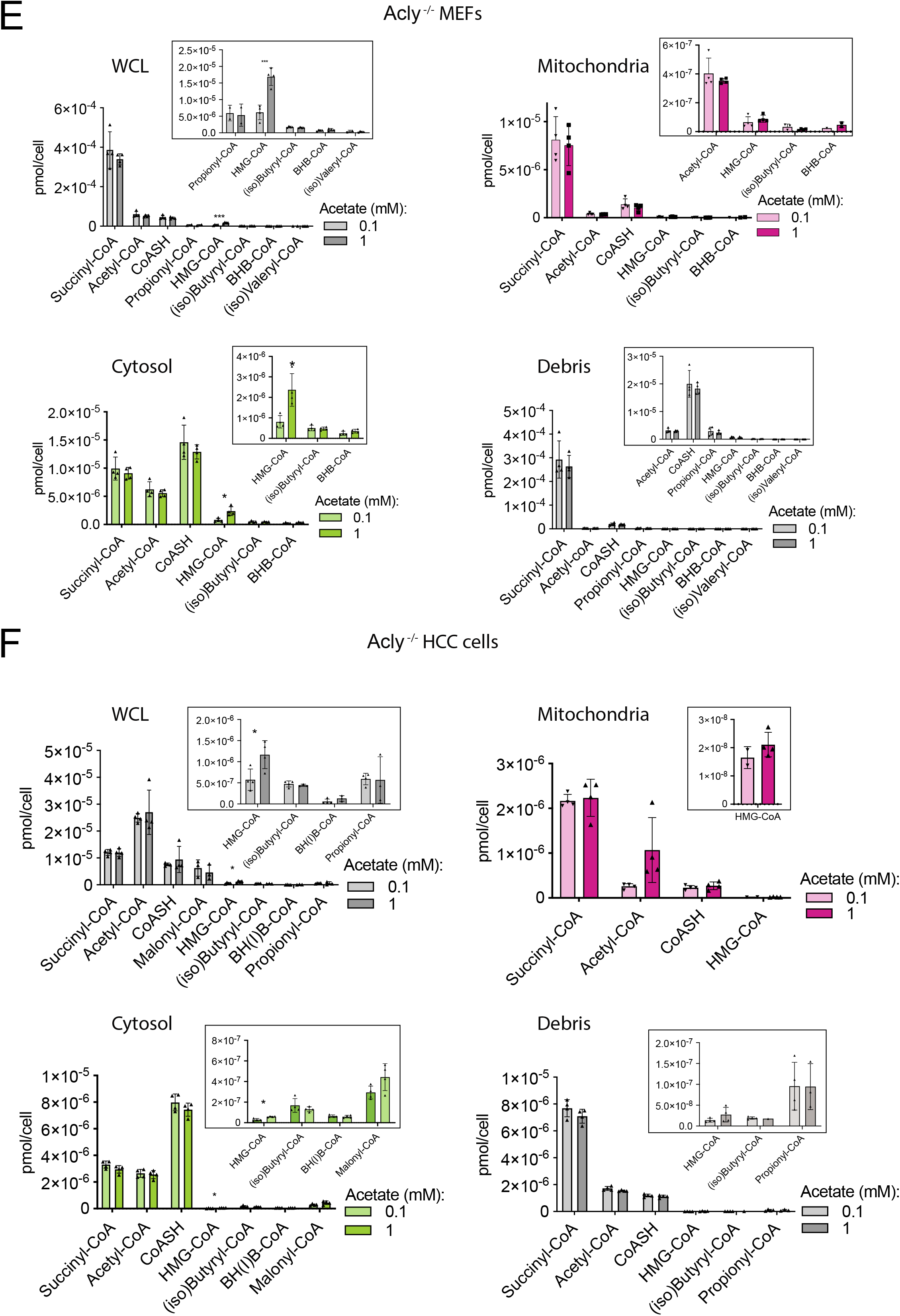
A) Western blot confirming ACLY deletion and ACSS2 upregulation in *Acly^-/-^* liver cancer cell line compared to control *Adly^f/f^* liver cancer cell line. Equal protein was loaded. B-F) Cells incubated in DMEM supplemented with 10% dialyzed fetal calf serum with the addition of the indicated concentration of acetate for 4 h. B-D) Data from Fig 3B, displayed as cellular acyl-CoA concentration. Metabolites were directly extracted from whole cells E,F) Representative experiments for SILEC-SF quantitation by differential centrifugation method. Data for all acyl-CoA species that were quantified in each fraction are displayed. Those that were not quantified showed insufficient signal intensity for the analyte, the internal standard or both. Low abundance metabolites are magnified in the upper right corner of each panel. Each symbol represents an individual replicate cell dish (n=4) from representative experiments. For all panels, error bars show standard deviation and statistical comparisons between two groups, were made by two-tailed Student’s ř-test with Welch’s correction and statistical significance defined as p < 0.05 (*), p < 0.01 (**), p < 0.001 (***), p < 0.0001 (****).

**Supplemental Figure 5.**
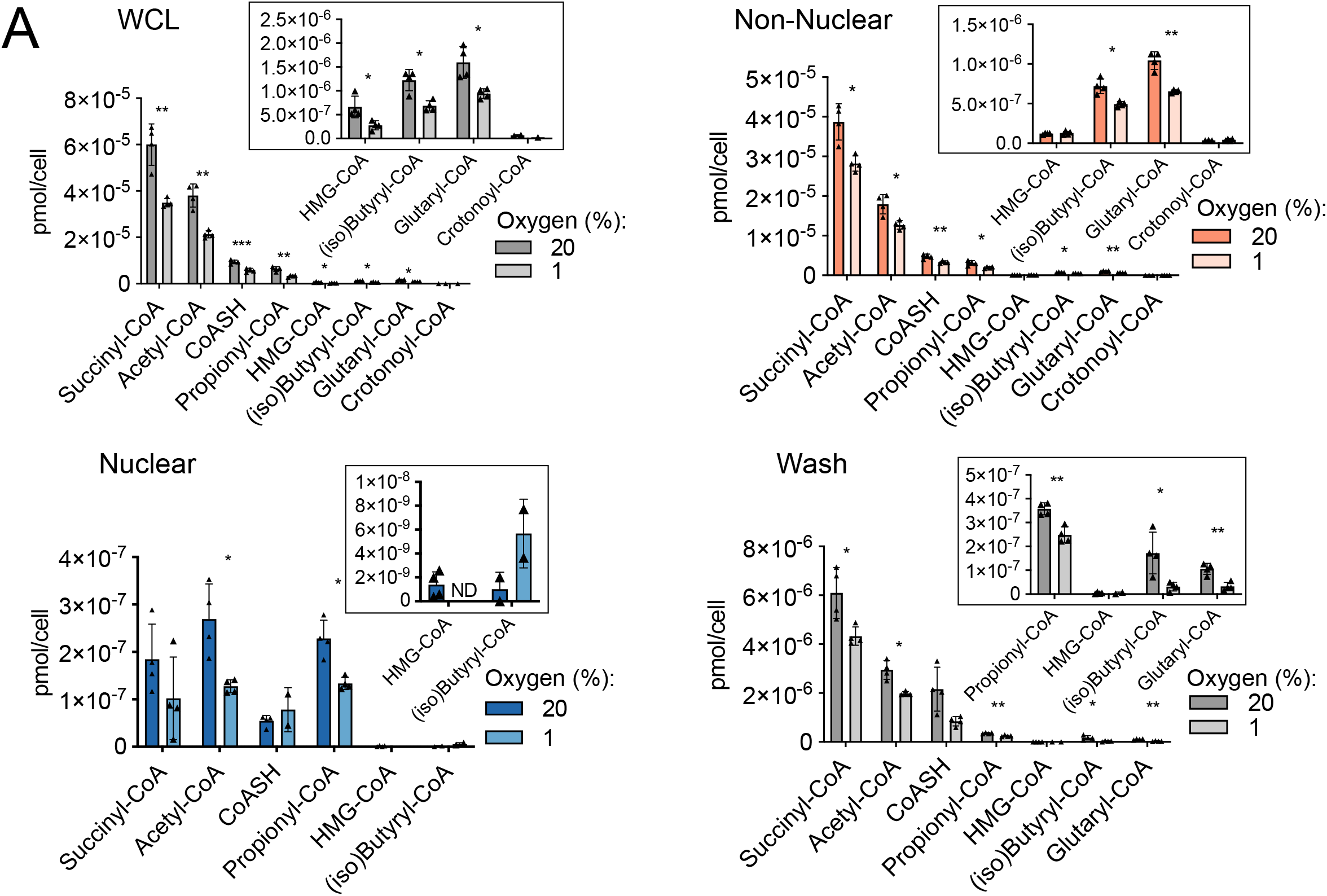
**A)** Data relating to **Fig 5D** and **5E.** HepG2 cells were incubated under 20% (normoxia) or 1% (hypoxia) for 24 h. Cells were serum starved in DMEM containing 5 mM glucose and 2 mM glutamine under their respective oxygen tensions for 2 h before harvest. All short chain acyl-CoA species quantified by SILEC-SF using the nuclear differential centrifugation procedure are shown from representative experiment in **Fig 5D** and incorporated into **Fig 5E.** Symbols represent individual replicate dishes (n=4) and error bars represent standard deviation. Lower abundance metabolites are magnified in the upper right corner of each panel. Error bars show standard deviation. Short chain acyl-CoA species that were not quantified showed insufficient signal intensity for the analyte, the internal standard or both. ND= not detected. Low abundance metabolites are magnified in the upper right corner of each panel. For comparison between two groups, datasets were analyzed by two-tailed Student’s ř-test with Welch’s correction and statistical significance defined as p < 0.05 (*), p < 0.01 (**).

**Supplemental Figure 6.**
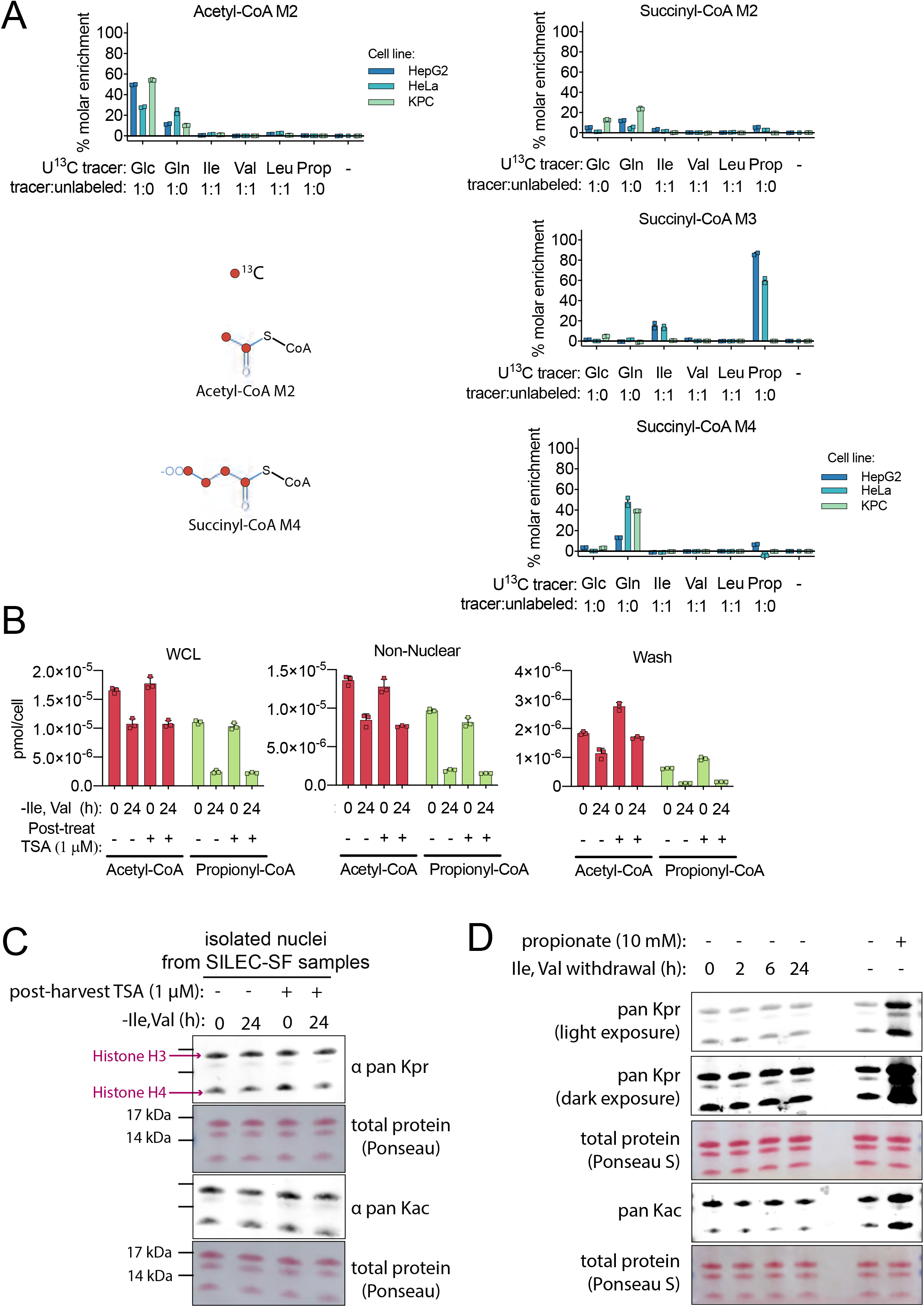
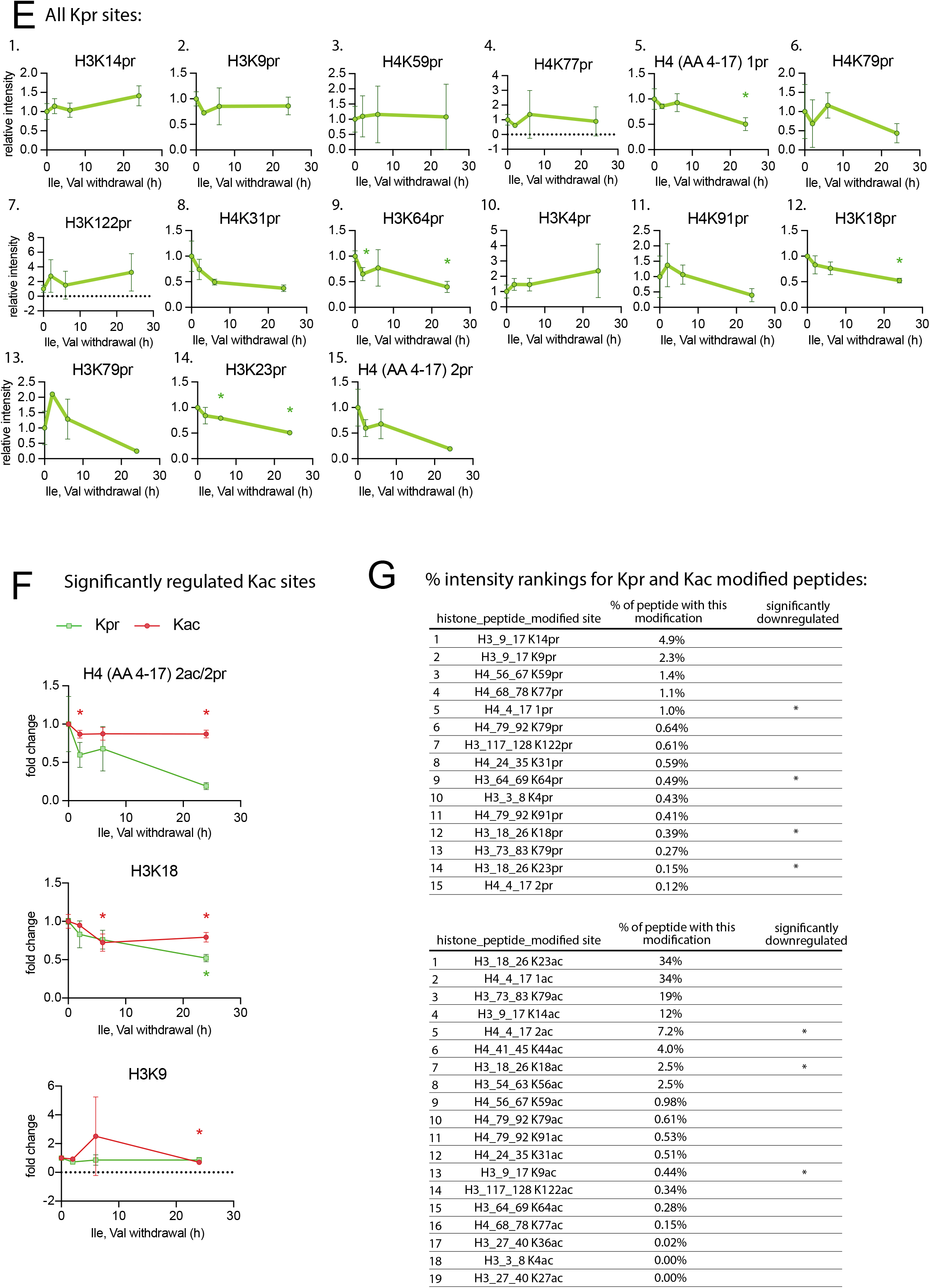
**A)** Incorporation of various substrates into acetyl-CoA and succinyl-CoA isotopologues was compared in whole cells by direct extraction of pancreatic adenocarcinoma cells (KPC) incubated in media containing uniformly (U) ^13^C-labeled substrates for 18 h. Total substrate concentrations were equal across all samples except for propionate, which was added only to the U^13^C_3_-propionate samples. U^13^C-labeled Val, Leu and Ile were diluted 1:1 with unlabeled substrate. N=3 replicate samples per condition from a single experiment. B,C) Data relating to **Fig 6G. B)** SILEC-SF analysis with nuclear/non-nuclear fractionation. Samples were treated with trichostatin A (TSA) in fractionation buffer. **C)** Western blots of protein pellets. **D)** Data relating to **Fig 6F-I.** Western blot of acid-extracted histones. **E)** relative intensity over time for all Kpr sites detected by acyl proteomic analysis. F) Relative intensity over time for 3 Kac sites identified as significantly regulated and for corresponding Kpr marks. G) Ranked average intensity across all timepoints of specific Kpr and Kac modified sites as a % of total signal detected across all modifications for that peptide. For all panels, error bars represent standard deviation. Statistical significance in **E & F** was determined by comparison to t=0 control for each mark and is indicated above or below the specific timepoint in the color matching the mark (p<0.05 (*).

